# Antigen-driven expansion of public clonal T cell populations in inflammatory bowel diseases

**DOI:** 10.1101/2024.05.15.594220

**Authors:** Mitchell Pesesky, Ramit Bharanikumar, Lionel Le Bourhis, Hesham ElAbd, Elisa Rosati, Cara L. Carty, Namita Singh, Bernd Bokemeyer, Stefan Schreiber, Siegfried Görg, Marco Garcia Noceda, Paidamoyo Chapfuwa, Rachel M. Gittelman, Damon May, Jennifer N. Dines, Wenyu Zhou, Ian M. Kaplan, Thomas M. Snyder, H. Jabran Zahid, Julia Greissl, Haiyin Chen-Harris, Bryan Howie, Andre Franke, Harlan S. Robins, Matthieu Allez

## Abstract

**Background:** Inflammatory Bowel Diseases (IBDs), including Crohn’s disease (CD) and ulcerative colitis (UC), are known to involve shifts in the T-cell repertoires of affected individuals. These include a reduction in regulatory T cells in both diseases, increase in TNFα production in CD, expansion of an unconventional T-cell population in CD, and clonal expansion of abundant T-cell populations in CD mucosal tissue. There are also differential HLA risk and protective alleles between CD and UC, implying CD- and UC-specific repertoire changes that have not yet been identified.

**Methods:** We performed ImmunoSequencing on blood samples from 3,853 CD cases, 1,803 UC cases, and 5,596 healthy controls. For each sample we imputed HLA type and cytomegalovirus (CMV) infection status based on public T-cell receptor β (TCRB) usage and identified public TCRBs enriched in CD or UC cases.

**Findings:** We determine that there is more expansion across clonotypes in CD, but not UC, compared with healthy controls. We also identify novel interactive effects of HLA-DQ heterodimers with CD and UC risk. Strikingly, from blood we identify public TCRBs specifically expanded in CD or UC. These sequences are more abundant in intestinal mucosal samples, form groups of similar CDR3 sequences, and can be associated to specific HLA alleles. Although the prevalence of these sequences is higher in ileal and ileocolonic CD than colonic CD or UC, the TCRB sequences themselves are shared across CD and not between CD and UC.

**Interpretation:** There are peptide antigens that commonly evoke immune reactions in IBD cases and rarely in non-IBD controls. These antigens differ between CD and UC. CD, particularly ileal CD, also seems to involve more substantial changes in clonal population structure than UC, compared to healthy controls.

## Introduction

### IBDs are prevalent and complex

Inflammatory bowel diseases (IBDs), including Crohn’s disease (CD) and ulcerative colitis (UC), are characterized by an immune-mediated uncontrolled inflammation of different segments in the digestive tract. IBDs affect over 2.3 million Americans and there are 70,000 new cases annually^1^. In Europe, between 2.5-3 million people are affected by IBDs^2^. Over the last two decades the rate of incidence of IBDs has increased in the Middle-East, Asia and South America^3^. IBDs are associated with multiple pathogenic factors, including environmental changes, an array of susceptibility gene variants, an abnormal microbiota composition, and exacerbated innate and adaptive immunity. There is an important heterogeneity among patients with both CD and UC, with diverse clinical presentations and phenotype, disease location, behavior and outcomes. While research in the last 10 years has advanced our understanding of these diseases and led to new therapeutic options, the exact mechanisms underlying these pathologies are still imperfectly understood – to this day, there is no curative treatment for patients.

### T cells are central to IBD pathogenesis

Modification of T-cell populations and functions, including an increase in CD4+ and CD8+ effector T cells and decrease in regulatory T cells are shown in the blood and intestinal mucosa of patients. T cell modulating molecules are used or in development for the treatment of IBDs; however, the antigens, cellular partners, and functional consequences of T cell reactivation in IBD patients’ mucosa remain to be fully understood. T cells are generated in the thymus through a tightly controlled process that generates T-cell receptors (TCRs) recognizing vast numbers of possible antigens in the context of <u>H</u>uman <u>L</u>eukocyte <u>A</u>ntigens (HLA) molecules.

During this differentiation, unique sequences at the TCR locus are created de novo to constitute a repertoire of possible receptors in a somatic recombination process known as V(D)J recombination^4,5^. A T cell’s reactivity towards an antigen leads to its proliferation and the generation of specific effector T-cell clones. These clones migrate into appropriate tissues to participate in local responses to their cognate antigen. Expanded clones populate the intestinal mucosa where they contribute to tissue homeostasis. The antigens driving these responses *in situ* could emanate from the microbiome present in the intestine.

HLA diversity impacts the naïve T-cell repertoire and could impact the immune responses in the context of diseases. Several MHC haplotypes, in both class I (HLA-A, -B and -C) and class II (HLA-DR, -DP, -DQ), have been associated with IBD. Some alleles are associated with both CD and UC as risk or protective modifications. Others are specifically associated with one form of IBD. As examples, *HLA-DRB1*07* is associated with ileal forms of CD, while *HLA-DRB1*15* is protective for CD but increases risk for UC^6^. Mechanistically, some variations in the HLA genes affect the peptide binding pocket of the molecule and the antigens that are presented to T cells. Other alleles impact other parts of the HLA molecules, potentially resulting in the selection of a different naive T-cell repertoire. Some alleles lead to variation in the amount of HLA complex at the surface of presenting cells, which would indirectly impact the T-cell response. Variation in the non-coding region of the HLA locus on chromosome 6 have also been associated with more aggressive disease phenotype in CD.

### T-cell repertoire analyses identify disease-associated populations

The TCR repertoire is a proxy to analyze antigen reactivity, and TCR sequencing can identify and classify patients with various infectious or inflammatory diseases, such as SARS-CoV2^7^ and Lyme disease^8^. Previous work has attempted to similarly identify specific T-cell clones which can differentiate these diseases^9–11^. These studies have not identified conventional HLA-binding TCR sequences associated with IBDs. Many of the cohorts of patients used are limited in numbers which could severely underpower the analyses, considering the tremendous diversity of the TCR repertoire. Nonetheless, certain features such as clonality and diversity were suggested to be modified in both blood and intestinal mucosa compared to controls and may differentiate CD and UC.

Studies have shown differences in T-cell population structure between CD and UC. Stimulating CD2 and CD8 resulted in increased IFN-γ secreting CD4+ T cells in CD but not in UC^12^. Additionally, studies have shown that Th1- and Th2-mediated responses are different in CD and UC^13^. One study also showed reduced TCR diversity in inflamed mucosa of patients with CD relative to healthy controls and patients with UC^14^.

Recently, mucosal T-cell clones were shown to be associated with disease recurrence after surgery in patients with ileal CD^15^. None of these expanded clones are common to different patients but the same TCR can be detected in samples from the same patient at different time points and locations, showing their spatial and temporal persistence. Interestingly, these clones are mostly found in the CD8 T-cell compartment and are associated with smoking, a common clinical feature of CD patients. More recently, specific clones with a semi-invariant TCR alpha chains have been shown to be expanded in CD patients. Upon further functional investigations these clones were shown to exhibit a distinct phenotype that resembles type II natural killer T cells as well as recognize antigens in the context of the invariant HLA-like molecule CD1d which presents lipids, glycolipids among others to T cells. These distinct features of these cells cause the author to propose them as a novel T-cell subset that was named (Crohn associated invariant T cells, CAIT)^10^. Despite their characterization, the antigen causing the expansion of these cells remains to be elucidated.

Although these studies have contributed to a better understanding of the role of T cells in IBD, several questions remain unanswered regarding the association of expanded or shared public clones with patient’s clinical history, disease progression and response to treatments. In this study based on samples collected in large international cohorts of IBD patients with different phenotypes and disease location, we show that the TCR repertoire displays increase clonality in IBD patients, independently of age and CMV infectious status. We identify previously uncharacterized public TCR sequences associated very specifically with CD or UC.

More precisely, these enhanced sequences (ES) were particularly abundant and characteristic of ileal CD compared to colonic CD and UC. HLA inference confirmed the association of some alleles with disease location and disease types, and identified new associations, such as DQ7.5 with CD. Finally, we show that groups of related ES associate robustly with certain HLAs advocating for the recognition of common peptides in patients with the same alleles.

## Methods

### Cohort Description

This study was based on samples collected in several observational study cohorts, totaling 3,853 CD cases, 1,803 UC cases, and 5,596 healthy controls. Blood samples were available in all patients. In addition, matched intestinal tissue samples from 450 of those individuals were also available. We randomly assigned blood samples to the sequence discovery or sequence validation sets to achieve an approximately 80:20 ratio, respectively (Table 1). Deidentified comparison case samples from other diseases were drawn from Adaptive’s internal databases to check cross-reactivity of CD- and UC-enriched TCRBs.

**Table 1:** Balance of samples in discovery and validation sets. IQR = interquartile range between 1^st^ and 3^rd^ quartile. N=number of samples with data annotated (all samples in category if not given).

| Region | Disease state | Sample type | Sequence Discovery (N) | Sequence Validation (N) | Mean age (IQR) | % Female | Mean years since diagnosis (IQR, N) |
| --- | --- | --- | --- | --- | --- | --- | --- |
| USA | Crohn's Disease | Peripheral Blood | 1496 | 341 | 41 (29-51) | 58% | 15 (7-20, 1366) |
| EU | Crohn's Disease | Peripheral Blood | 1866 | 384 | 38 (26-47) | 60% | 9 (1-14, 1159) |
| USA | Ulcerative colitis | Peripheral Blood | 774 | 195 | 43 (31-54) | 50% | 12 (4-18, 706) |
| EU | Ulcerative colitis | Peripheral Blood | 669 | 165 | 42 (30-52) | 50% | 29 (23-34, 55) |
| USA | Healthy | Peripheral Blood | 1874 | 454 | 38 (28-47) | 52% | NA |
| EU | Healthy | Peripheral Blood | 2694 | 574 | 50 (37-60) | 47% | NA |
| EU | Crohn's Disease | Intestinal Mucosa | 0 | 406 | 36 (26-43) | 56% | 10 (3-14, 406) |
| EU | Ulcerative colitis | Intestinal Mucosa | 0 | 44 | 42 (33-50) | 41% | 10 (3-17, 42) |
| EU | MS | Peripheral Blood | 0 | 1736 |  |  |  |
| USA | MS | Peripheral Blood | 0 | 636 |  |  |  |
| EU | RA | Peripheral Blood | 0 | 1219 |  |  |  |
| USA | Celiac | Peripheral Blood | 0 | 923 |  |  |  |
| EU | COVID | Peripheral Blood | 0 | 429 |  |  |  |
| USA | COVID | Peripheral Blood | 0 | 21 |  |  |  |
| EU | IBS | Peripheral Blood | 0 | 390 |  |  |  |
| USA | Lyme | Peripheral Blood | 0 | 216 |  |  |  |
| USA | CRC | Intestinal Biopsy | 0 | 2169 |  |  |  |

### Immunosequencing

Genomic DNA was extracted from frozen, plasma-depleted blood samples using the Qiagen DNeasy Blood Extraction Kit (Qiagen). As much as 18 μg of input DNA was then used to perform immunosequencing of the third complementarity determining regions (CDR3) of TCRB chains using the ImmunoSEQ Assay (Adaptive Biotechnologies). Briefly, input DNA was amplified in a bias-controlled multiplex PCR, followed by high-throughput sequencing. Sequences were collapsed and filtered to identify and quantitate the absolute abundance of each unique TCRB CDR3 region for further analysis, as previously described^7,16,17^ In order to quantify the proportion of T cells out of total nucleated cells input for sequencing, or T-cell fraction, a panel of reference genes present in all nucleated cells was amplified simultaneously^18^.

### Clone size distribution metrics

All repertoires were stochastically downsampled to 50,000 total TCR read templates. Clonotype count (R) was the total number of clonotypes in the sample after downsampling. Simpson clonality was calculated as 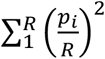 where p is the number of templates in the i-th largest clonal group. Singleton breadth was calculated as 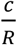 where c was the number of clonal populations with only a single template in the sample. Distributions of these metrics were compared between disease states (CD, UC, or healthy control) using the Kolmogorov–Smirnov (KS) test, against the null hypothesis that both conditions were sampling the same underlying distribution.

### Enhanced sequence discovery

Identification of CD- or UC-specific TCRBs, also known as <u>e</u>nhanced <u>s</u>equences (ES), was performed as described previously^7^. Briefly, we assessed occurrence in disease cases and healthy controls for each TCRB in the sequence discovery set and determined statistical enrichment in cases using a one-sided Fisher’s Exact Test (FET). TCRBs with a FET p-value below 0.001 (threshold determined empirically) were added to the ES set.

### HLA association and interaction effects

HLA types for each individual in the sequence discovery set were inferred from the T-cell repertoire as described previously^19^. Briefly, we identified TCRs specific to each of the 145 most common HLAs in a separate population from the United States, using the same ES discovery method described above to identify HLA-specific ES instead of disease-specific ES. We then used those HLA-specific ES to build regression-based HLA classifiers, and applied those classifiers to the TCR repertoires in the CD- or UC-ES discovery set. The classifier results were then used as the imputed HLA type for that sample. Individual ES were associated to an HLA by identifying the HLA with the lowest FET p-value for cooccurrence with the ES in the sequence discovery set. The association was considered confident if this p-value was less than 1×10^−4^.

HLA enrichment in patients compared to controls was assessed for single and combination alleles using the Fisher’s Exact Test (FET), with correction for multiple comparison. Odd’s ratio (OR) was used to identify additive, non-additive and interaction effects at the HLA loci in CD relative to healthy controls.

### Enhanced sequence clustering

ES were encoded as a CDR3 amino acid sequence, V-gene token, and J-gene token. These encodings were then represented as nodes in a network, where nodes were connected by an edge if they had matching V and J genes, and their CDR3 had a single amino acid mismatch (1-Hamming distance). Connected component groups, consisting of all ES that are connected through any number of nodes and edges are then referred to as ES clusters.

## Results

### Impact of IBD on clonotype size distribution: evidence of peripheral clonal T-cell expansions in CD and UC

Our aim was to identify specific patterns in the repertoire of T-cell receptor (TCR) sequences that can serve as a biomarker for IBD. To ensure that we could test any identified biomarkers on independent samples, we randomly separated our data, with 80% going into a ‘sequence discovery’ set and 20% into a ‘validation’ cohort. The sequence discovery cohort included peripheral blood samples from 3,128 CD cases, 1,443 UC cases, and 4,568 healthy control (HC) individuals. Only 21 control samples had fewer than 50,000 clonotypes sequenced (Fig S1A) so, to better balance the data, we removed all samples below that threshold from future analysis. This left 2,975 CD cases, 1,308 UC cases, and 4,549 healthy controls (Table 1). The distribution of unique clonotypes per sample was significantly shifted downwards for IBD cases relative to healthy controls (p < 1×10^−200^, KS test, Table 2, Figure S1A). To determine if this difference reflected DNA extraction batch effects or an IBD-driven decrease in T-cell abundance, we also checked the T-cell fraction of the total cell DNA derived from each sample. These values were also significantly decreased (Figure S1B, p < 1×10^−100^, KS test). Unique clonotypes and productive T-cell fraction were also significantly decreased in UC cases compared to healthy controls (p < 1×10^−100^ and p < 1×10^−45^, respectively, KS test), though with less clear separation than the CD samples. There was no significant difference between CD and UC in the total clonotypes sequenced, and productive T-cell fraction was significantly lower in CD as compared to UC (p = 0.003).

**Table 2:** Sequence discovery cohort characteristics.

| Diagnosis | Samples passing QC | Median clonotypes | Median % T cells |
| --- | --- | --- | --- |
| CD | 2,975 | 159,527 | 18% |
| UC | 1,308 | 158,334 | 19% |
| HC | 4,549 | 266,788 | 24% |

We then assessed the effect of IBD on the clone size distribution of the T-cell repertoire. Prior work has demonstrated substantial differences in clone size distribution depending on an individual’s age and CMV status. We restricted our analysis to individuals between 20 and 79 years old and binned them into three 20 year age bins. We also used our previously published CMV classifier^20^ to separate individuals based on CMV status. Because of our large starting set, each group of samples represents at least 100 individuals, even after splitting by diagnosis, age bin, and CMV status. We then downsampled each repertoire to 50,000 read templates to allow comparisons between samples and then applied three metrics to quantify shifts in clonotype size distribution.

The first metric, Simpson’s Clonality, is a measure of how even the clonotype population sizes are, and differences are driven primarily by differences in large clonal populations (Figure 1A). The second metric, clonotype count, represents the total diversity of clonotypes in the sample. Differences in this metric can be driven by clonotypes of any size (Figure 1B). The third metric, singleton breadth, measures the proportion of clonal populations that have the lowest level of expansion observed, and it is driven by changes in smaller clonal populations (Figure 1C).

**Figure 1:**
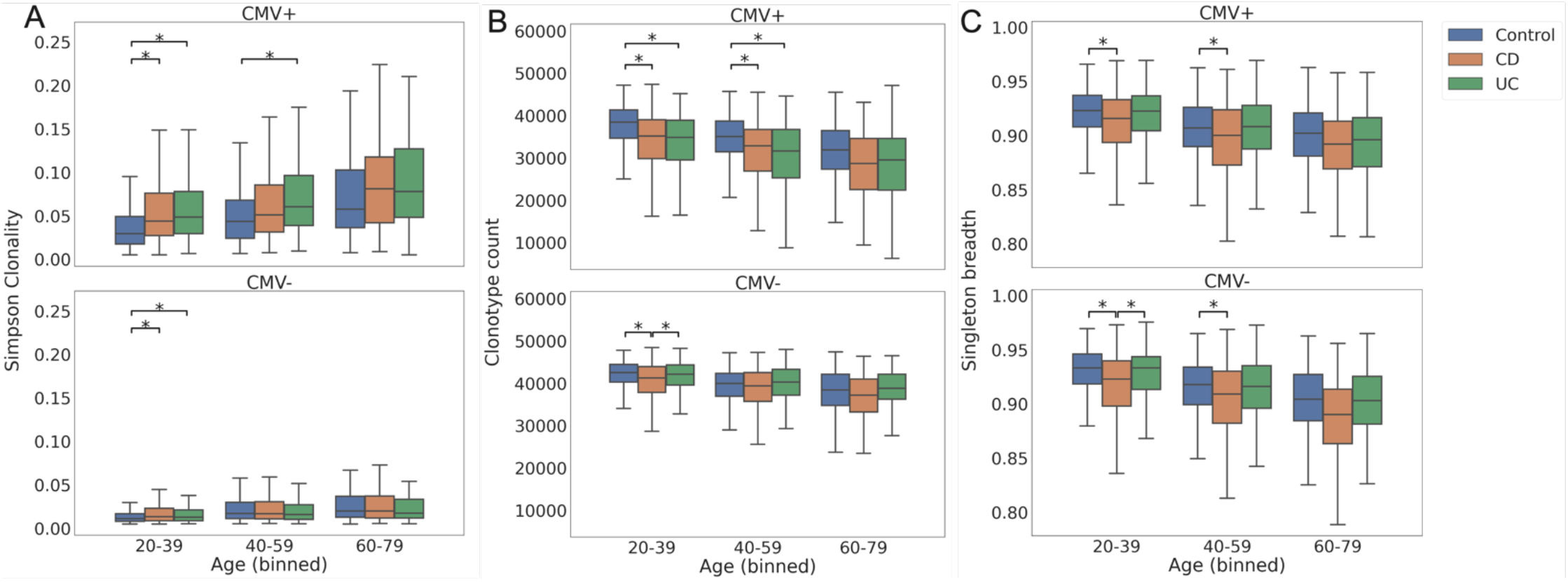
After downsampling to 50,000 total read templates in all samples from individuals with attached age data, we assessed A) Simpson clonality, B) clonotype count, and C) singleton breadth for CD and UC cases and healthy controls, across age bins and CMV status. All pairwise comparisons within an age and CMV bin were assessed with a Kolmogorov–Smirnov test for equivalence of the distributions. All comparisons with p < 0.05 are indicated with a ‘*’, though in this case all marked comparisons have p < 2×10^−4^. Each boxplot represents the distribution of normalized clonotype counts in that bin across samples. Boxes cover the first and third quartile of the distribution (with the median indicated within the box), while whiskers show the remainder of the distribution up to the interquartile range. Outliers were excluded to better visualize the central quantiles of the distribution.

Using these three metrics, we see that CD and UC cases have elevated clonality compared to healthy controls in CMV+ individuals and the differences are statistically significant (p < 3×10^−4^) in individuals aged 20-39. We also observe that CD and UC cases have reduced downsampled clonotype counts in CMV+ individuals, but in CMV-individuals only CD cases show reduced downsampled clonotype counts. Finally, we observe that CD, but not UC, has reduced singleton breadth compared to healthy controls, across age bins and CMV status. These measures suggest two different patterns of expansion. First, in CMV+ individuals, the largest clonal populations in CD and UC cases are significantly larger than in heathy controls, as seen by differences in clonality (Figure 1A, Table S1). The second pattern is a broad expansion of clonal populations in CD cases. This is particularly evident in its effect on singleton breadth which is significantly lower than in control samples in all demographics except CMV+ ages 60-79 (Figure 1C). The combination of these two patterns is a reduction in sampled diversity in CD and UC, particularly in CMV+ individuals, but also for CD in CMV-individuals age 20-39 (Figure 1B). These patterns point to some potentially interesting biological differences in the adaptive immune response to CD and UC.

### Imputed functional HLA types expand our knowledge of HLA interaction with IBDs

To complement our investigation of the T-cell repertoire, we also explored interaction effects of individual HLA alleles with CD and UC. Due to the multi-study nature of our cohort, HLA typing was not completed for all samples. Leveraging recently developed models for determining HLA type from TCR repertoires, we wished to examine HLA interactions with IBD risk from a T-cell perspective. The T-cell perspective differs from a genetic perspective of HLA interaction in that it emphasizes functional HLA complexes rather than individual genes. Practically, this means increased focus on DP and DQ heterodimers, and decreased focus on HLA alleles with low surface expression, particularly HLA-C.

We confirm here previously reported risk/protective effects for different HLA alleles in CD^6,21^. Under an additive model, we found a significant risk effect for DRB1*07:01 and protective effects for DRB1*04:01 and DRB1*15:01 (Table 3) in Crohn’s cases relative to controls. Additionally, we also identified novel zygosity and interaction effects at the HLA DQ locus. Specifically, DQ7.5 provides a significant CD risk, when combined with DQ2.2 or DQ4.4, potentially indicating the formation of trans-heterodimers of those alleles. In contrast, DQ6.2 provides a strong protective effect regardless of zygosity.

**Table 3.** Zygosity and interactions effects at the HLA Class II loci in Crohn’s disease compared to healthy controls.

| Genotype | Reference | OR [95% CI] | p-value |
| --- | --- | --- | --- |
| DRB1*07:01/DRB1*07:01 | DRB1*X/DRB1*X | 1.64[1.13-2.39] | <0.001 |
| DRB1*07:01/DRB1*07:01 | DRB1*07:01/DRB1*X | 1.97 [1.39-2.81] | <0.001 |
| DRB1*15:01/DRB1*15:01 | DRB1*X/DRB1*X | 0.51 [0.34-0.78] | <0.001 |
| DRB1*04:01/DRB1*04:01 | DRB1*X/DRB1*X | 0.54 [0.30-0.96] | 0.03 |
| DQ7.5/DQ2.2 | DQX/DQY | 1.84 [1.32-2.58] | <0.0001 |
| DQ7.5/DQ4.4 | DQX/DQY | 2.28 [1.26-4.49] | <0.001 |
| DQ6.2/DQ6.2 | DQX/DQX | 0.46 [0.32-0.65] | <0.0001 |
| DQ6.2/DQ6.2 | DQ6.2/DQY | 0.69 [0.50-0.95] | 0.02 |

We also observed certain HLA alleles conferring differential risk in CD and UC. While DRB1*15:01 was protective in CD, it was associated with a non-additive risk effect (OR = 2.18, 95% CI = [1.50-3.19]) in UC. The opposite trend was observed with DRB1*07, which had a risk effect in CD and a dominant protective effect in UC (OR = 0.50, 95% CI = [0.30-0.85]). We also observed increased rates of HLA Class 2 Homozygosity in UC compared to both Crohn’s and healthy controls. These results suggest the involvement of different clonotypes and antigens in the two diseases.

### Specific, public TCRB sequences associate with CD and UC in peripheral blood

We next sought to determine if there were any specific TCRBs enriched in CD and UC. For each TCRB occurring in more than one individual in our set, we performed a one-sided Fisher’s Exact Test (FET) to determine if these TCRB sequences were significantly enriched in IBD cases over healthy controls. We defined Enhanced Sequences (ES) for a disease of interest to be those TCRB sequences that are highly enriched in patients compared to controls (citation of previous papers). At the threshold of p< 0.001, we discovered 1,883 CD ES and 155 UC ES. We observed no TCRB sequences in common between the two ES sets.

### IBD ES are significantly enriched in validation cases over healthy controls

To determine if the enrichment of IBD ES in IBD cases generalized to our Sequence Validation set, we needed to account for the observed difference in clonotype counts between cases and controls. Accordingly, we used the metric of sequence breadth to assess ES abundance in individual repertoires (Table 1), where breadth equals the count of ES divided by the count of total unique TCRB sequences in the sample. The CD ES occur at a significantly higher breadth in CD cases (mean breadth of 1.5 × 10^−4^, ie 1.5 ES per 10,000 clonotypes) than healthy controls or other diseases (Figure 2A). 46% of CD cases had a CD ES breadth in blood greater than the 99^th^ percentile of healthy controls (Figure S3).

**Figure 2:**
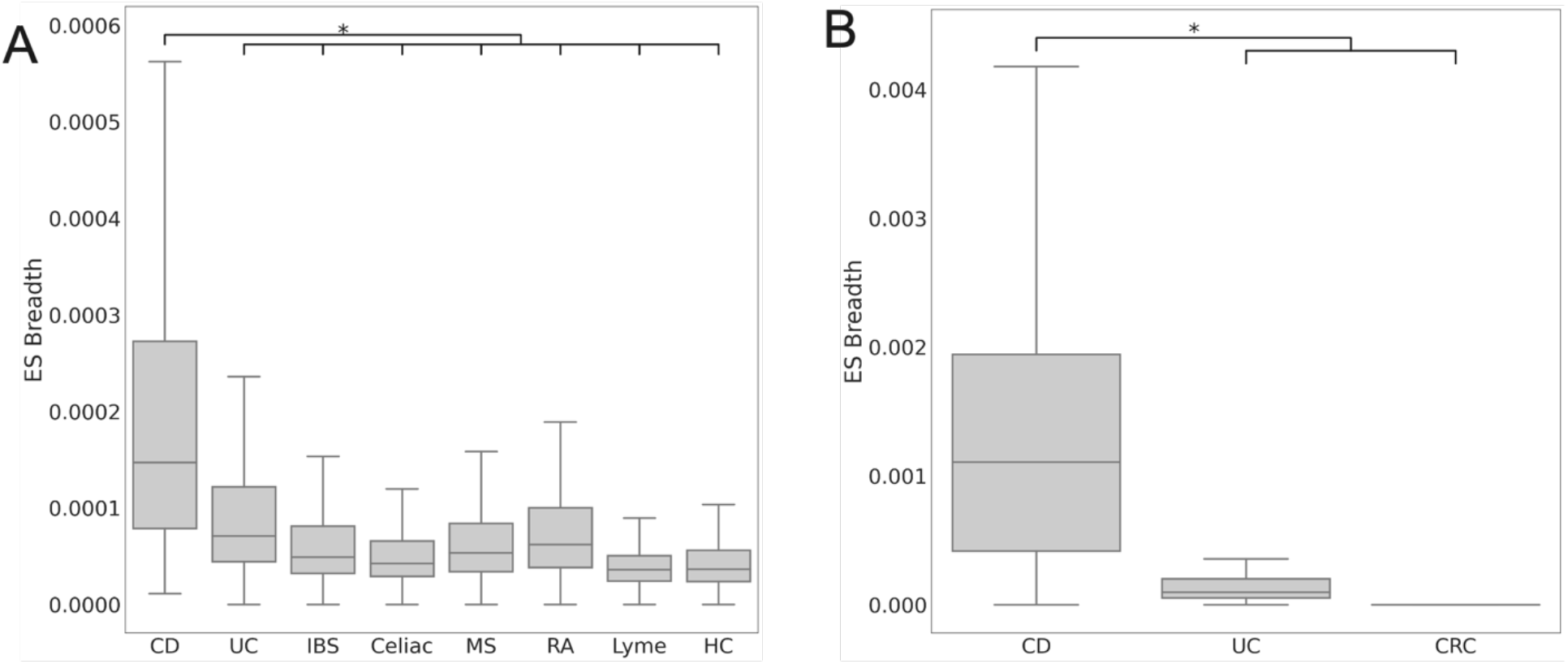
Distribution of sequence breadth of Crohn’s associated ES in (A) sequence validation blood samples and (B) tissue samples. Boxplots show first and third quartile values of the breadth range, with the median marked within the box. Whiskers extend to furthest point within 1.5 times the interquartile range. Points outside of the whiskers were excluded for visualization purposes. CD = Crohn’s disease, UC = ulcerative colitis, HC = healthy control, MS = multiple sclerosis, RA = rheumatoid arthritis, CRC = colorectal cancer. *p-value < 1×10^- 30^, Mann-Whitney U test.

We also looked at T cells sampled from intestinal mucosa from patients with CD, UC, or colorectal cancer (CRC), and found that the median CD mucosa sample had an order of magnitude higher CD ES breadth than the median CD blood sample, and that there was more complete separation between CD and non-CD mucosa samples than between CD and non-CD blood samples (Figure 2B).

UC ES have a lower median breadth across samples (averaged 5×10^−6^), and a lower difference in median breadth between UC samples and healthy controls. Nevertheless, the difference in UC ES breadth between UC cases and healthy controls is significant (Figure 3A). Consistent with our observations of CD ES, UC ES are 10 fold enriched in UC mucosal samples compared to blood, with a larger breadth difference between UC and CD than in blood samples (Figure 3B). These data suggest disease-associated TCRB sequences are highly enriched in patient blood compared to healthy controls and are found at an even higher concentration at the site of disease compared to circulation in both CD and UC patients.

**Figure 3:**
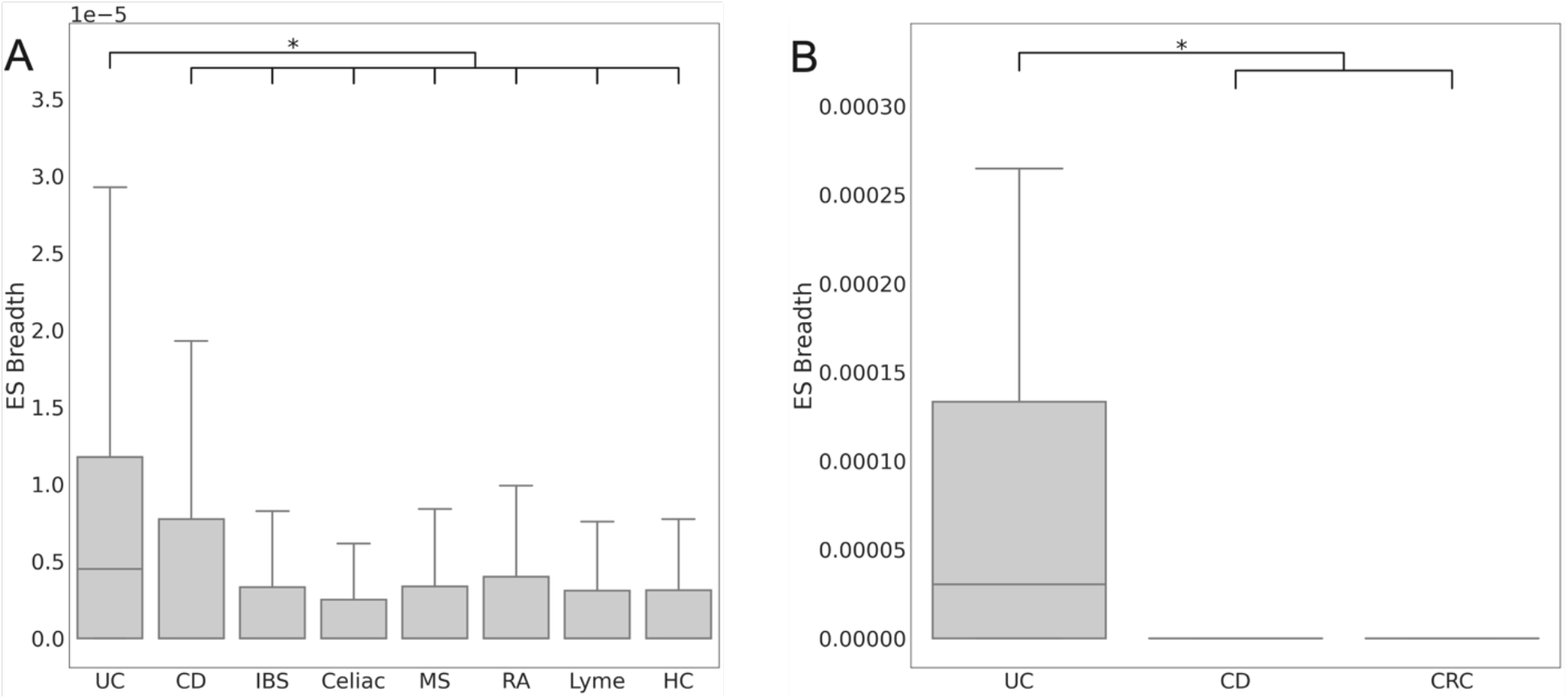
Distribution of sequence breadth of UC associated ES in (A) holdout blood samples and (B) holdout tissue samples. Boxplots show first and third quartile values of the breadth range, with the median marked within the box. Whiskers extend to furthest point within 1.5 times the interquartile range. Points outside of the whiskers were excluded for visualization purposes. CD = Crohn’s disease, UC = ulcerative colitis, HC = healthy control, MS = multiple sclerosis, RA = rheumatoid arthritis, CRC = colorectal cancer. *p-value < 1×10^−30^, Mann-Whitney U test.

### Some IBD ES observed in prior study of yeast-activated T cells

To evaluate whether our IBD ES sequences are novel, we performed a comprehensive search for the TCRB CDR3 amino acid sequences from the CD and UC ES in three public databases (VDJdb, McPAS, and IEDB). There was no overlap between these databases and the CD ES, but two UC ES were found in IEDB: the CDR3 CASSLELPPGETQYF matching the LSPRWYFYYL peptide from SARS-CoV2 and the CDR3 CASSLEGPGANVLTF matching the peptide GDAALALLLLDRLNQL also from SARS-CoV2. We also checked the CD ES against three recently published sets of CD-relevant TCR sequences. In the first two – the CAIT paired TCRB sequences reported by Rosati et al^10^, and the OmpC-reactive TCRs from Uchida et al ^11^– there is no overlap, but in a single cell analysis of T cells activated by yeast lysates in Crohn’s patients^22^ we observe an overlap of 26 TCRBs with the CD ES (Table 4). This suggest that a subset of the CD ES, but not the UC ES, can be activated by yeast derived antigens.

**Table 4:** TCRBs in common between CD ES and yeast activated T cells with the activation conditions in Martini et al. 2023.

| CDR3 | V gene | J gene | Martini et al. Condition |
| --- | --- | --- | --- |
| CASSLAPTVSYNEQFF | TCRBV05-01 | TCRBJ02-01 | S.cerevisiae-reactive CD4 T |
| CASSLELQETQYF | TCRBV07-03 | TCRBJ02-05 | C.tropicalis-reactive CD4 T |
| CASSLASGSRQETQYF | TCRBV05-04 | TCRBJ02-05 | C.tropicalis-reactive CD4 T |
| CASSLASGSRQETQYF | TCRBV05-04 | TCRBJ02-05 | S.cerevisiae-reactive CD4 T |
| CASSLDGLKETQYF | TCRBV11-01 | TCRBJ02-05 | S.cerevisiae-reactive CD4 T |
| CASSPAGGRNEQFF | TCRBV11-02 | TCRBJ02-01 | S.cerevisiae-reactive CD4 T |
| CASSRGASGNTIYF | TCRBV05-01 | TCRBJ01-03 | C.tropicalis-reactive CD4 T |
| CASSPLAGGPRTDTQYF | TCRBV11-02 | TCRBJ02-03 | S.cerevisiae-reactive CD4 T |
| CASSPLAGGPRTDTQYF | TCRBV11-02 | TCRBJ02-03 | C.tropicalis-reactive CD4 T |
| CASSSGGGRDNEQFF | TCRBV05-01 | TCRBJ02-01 | C.albicans-reactive CD4 T |
| CASSSTGTANQPQHF | TCRBV06-05 | TCRBJ01-05 | S.cerevisiae-reactive CD4 T |
| CASSSRHEQYF | TCRBV04-02 | TCRBJ02-07 | C.tropicalis-reactive CD4 T |
| CASSRLVGDTQYF | TCRBV07-02 | TCRBJ02-03 | C.tropicalis-reactive CD4 T |
| CASSYSPTGATEAFF | TCRBV06-05 | TCRBJ01-01 | S.cerevisiae-reactive CD4 T |

| **CDR3** | **V gene** | **J gene** | **Martini et al. Condition** |
| --- | --- | --- | --- |
| CASSPGQISNTEAFF | TCRBV07-02 | TCRBJ01-01 | C.tropicalis-reactive CD4 T |
| CASSPGQISNTEAFF | TCRBV07-02 | TCRBJ01-01 | S.cerevisiae-reactive CD4 T |
| CASSLELQETQYF | TCRBV07-02 | TCRBJ02-05 | C.albicans-reactive CD4 T |
| CASSLFGTGPYEQYF | TCRBV05-06 | TCRBJ02-07 | S.cerevisiae-reactive CD4 T |
| CSSNTDTQYF | TCRBV29-01 | TCRBJ02-03 | C.tropicalis-reactive CD4 T |
| CAISLEGGGSYEQYF | TCRBV10-03 | TCRBJ02-07 | S.cerevisiae-reactive CD4 T |
| CASSEERSGNTIYF | TCRBV25-01 | TCRBJ01-03 | C.albicans-reactive CD4 T |
| CASSLPGFSSYEQYF | TCRBV05-06 | TCRBJ02-07 | S.cerevisiae-reactive CD4 T |
| CASSLYGTGPYEQYF | TCRBV05-06 | TCRBJ02-07 | S.cerevisiae-reactive CD4 T |
| CASSASGTANQPQHF | TCRBV06-05 | TCRBJ01-05 | S.cerevisiae-reactive CD4 T |
| CASSLGGDSSYNEQFF | TCRBV07-02 | TCRBJ02-01 | C.tropicalis-reactive CD4 T |
| CASSLLGVSYEQYF | TCRBV05-06 | TCRBJ02-07 | S.cerevisiae-reactive CD4 T |
| CASKREIADTQYF | TCRBV05-01 | TCRBJ02-03 | S.cerevisiae-reactive CD4 T |
| CSASLTSGRAADTQYF | TCRBV20-01 | TCRBJ02-03 | S.cerevisiae-reactive CD4 T |
| CASSSENSPLHF | TCRBV06-06 | TCRBJ01-06 | S.cerevisiae-reactive CD4 T |

### More ES are found in ileal than colonic forms of IBD

Our initial ES discovery identified an order of magnitude more CD ES than UC ES. Possible factors underlying this finding are (1) sample size imbalance: we had 2.3 times more CD samples than UC samples, hence more statistical power to identify CD ES; (2) differences in T cells’ roles in CD and UC biology. Within CD, we observed that samples from patients with ileal involvement had higher CD ES breadth than those from patients with only colonic involvement (p < 1×10^−5^, Mann Whitney U test). Further, samples from patients with complications such as stricturing and/or fistulizing behavior showed higher trending ES breadth than samples without complications (Figure S5).

To eliminate the effect of sample size imbalance and to determine any association between ES abundance and disease characteristics we randomly sampled 500 repertoires each from our ileal only CD cases (L1), colonic only CD cases (L2), ileocolonic CD cases (L3), and UC cases. We repeated that sampling 10 times for each disease subtype to understand the variance in subsequent measurements. For each set of repertoires, we again identified enriched sequences using the FET. In this case, we generated lists with three different levels of stringency based on maximum p-value (10^−3^, 10^−4^, and 10^−5^).

This method demonstrated that at low p-values (<10^−5^) there are significantly (p<0.001) and substantially more TCRB sequences enriched in L1 and L3 samples than L2 or UC samples (Figure 4A). We saw no significant difference in number of enriched sequences between L2 and UC. Comparing the identity of TCRBs enriched in L1, L2, L3, and UC, we noted the highest overlap between L1 and L3 (Figure 4B). L2 has a significantly higher overlap of specific TCRs with L1 or L3 than with UC (Table S2). In fact, very few TCRs overlap between any of the CD subtypes and UC. These trends were similar at the three p-value thresholds tested, though L3 had the lowest median number of specific TCRs at p<10^−3^ (Figure S6). Aligned with these trends, we saw no overlap in ES between CD and UC from the fully powered sequence discovery. Taken together, these results demonstrate that the abundance of IBD-specific TCRs is highest in ileal disease cases, but CD ES are distinct from UC ES, regardless of disease location.

**Figure 4:**
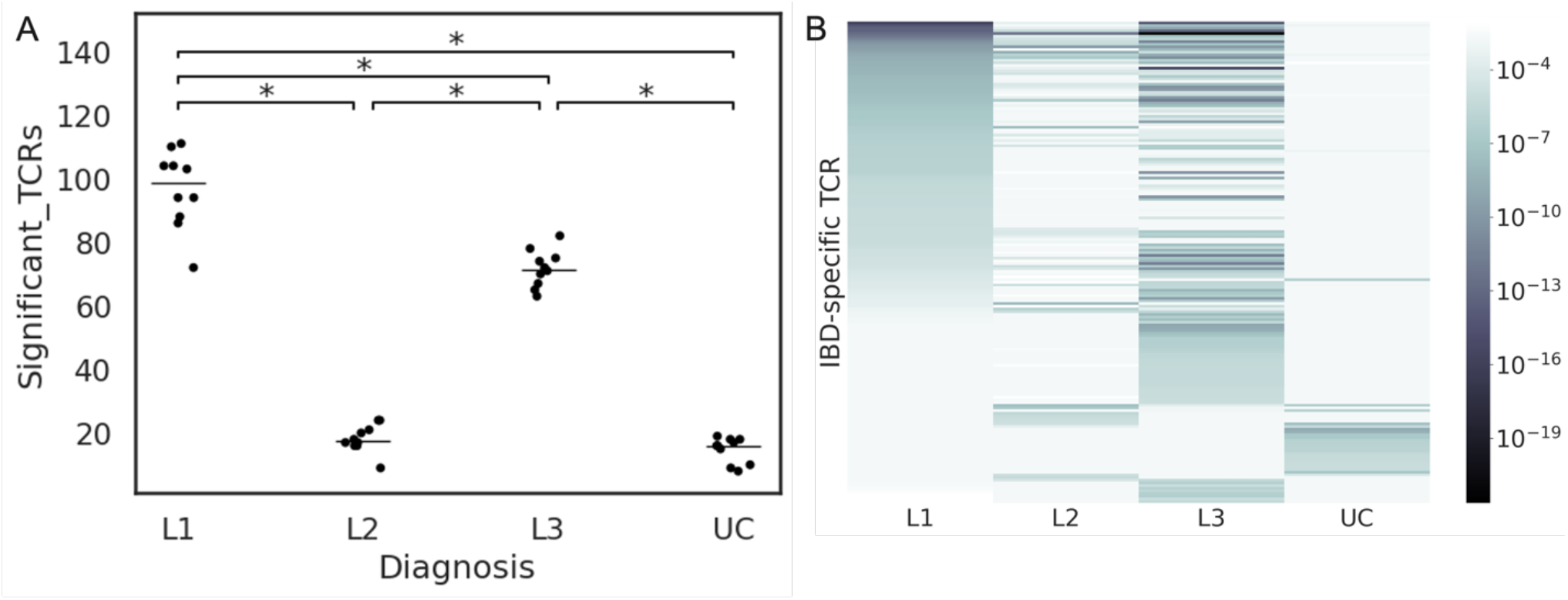
Identification of specific TRB sequences enriched in CD and UC. A) Count of TRB sequences in repeat experiments. Each dot represents an experiment comparing 500 randomly selected disease repertoires to 2,607 control repertoires. 10 experiments were performed for each disease. Median values shown as a horizontal line. *p<0.001. B) Heatmap of Fisher’s Exact Test (FET) p-values for TCRs associated to IBD subtypes from a single downsampling experiment. The y-axis is any TCR that had a p-value < 10^−5^ in at least one subtype, while the color gradient indicates it’s exact p-value for the subtype on the x-axis. TCRs are ordered vertically first by L1 p-value, then (in case of ties) L3, then L2, and finally UC. L1=ileal CD, L2=colonic CD, L3=ileocolonic.

### Individual high confidence CD ES are common in CD cases

We focused our investigation of TCRB characteristics on the CD ES since they more strongly separated disease cases from healthy controls. We further enriched the CD ES list for those T cells most likely to be involved in the disease by filtering the list with two criteria: 1) the ES was found at higher breadth in mucosa than blood, and 2) the ES was not one of the sequences that are a part of our HLA models (Figure S7). Our rationale for these filters was that TCRBs relevant and specific to the disease should be abundant in disease mucosa and too rare in healthy controls to build HLA models. This left 1007 high-confidence CD ES.

To aid in understanding these high-confidence ES, we evaluated more interpretable metrics of their prevalence in cases and controls. The distribution of high-confidence ES per 100,000 clonotypes is significantly lower for healthy controls than for CD cases (p<10^−197^), with 2.4% of CD cases exceeding 40 ES per 100,000 clonotypes (Figure S8A). The 10 ES with the lowest p-values are found in 3%-9% of CD cases and 0.1%-1% of healthy controls (Figure S8B).

### Most high confidence CD ES associate to class II HLAs

We associated each of the high-confidence ES to the HLA that best explained its presence or absence across samples, and used an FET to determine a p-value for the null hypothesis that the association was due to random co-variation. Using a strict p-value threshold of p<10^−4^, we determined that the HLA associated to the most ES was DR7 (DRB1*07:01), but that several other HLAs had nearly as many associated ES (Figure 5A). 738 (73%) of the high confidence ES were confidently associated to an HLA. Of those with confident associations, 676 (83%) were associated to class II HLAs (Figure S9). ES were associated to HLAs that were both over- and under-represented in disease cases compared to controls, and ES association was not strictly a function of overall HLA prevalence (Figure 5B).

**Figure 5:**
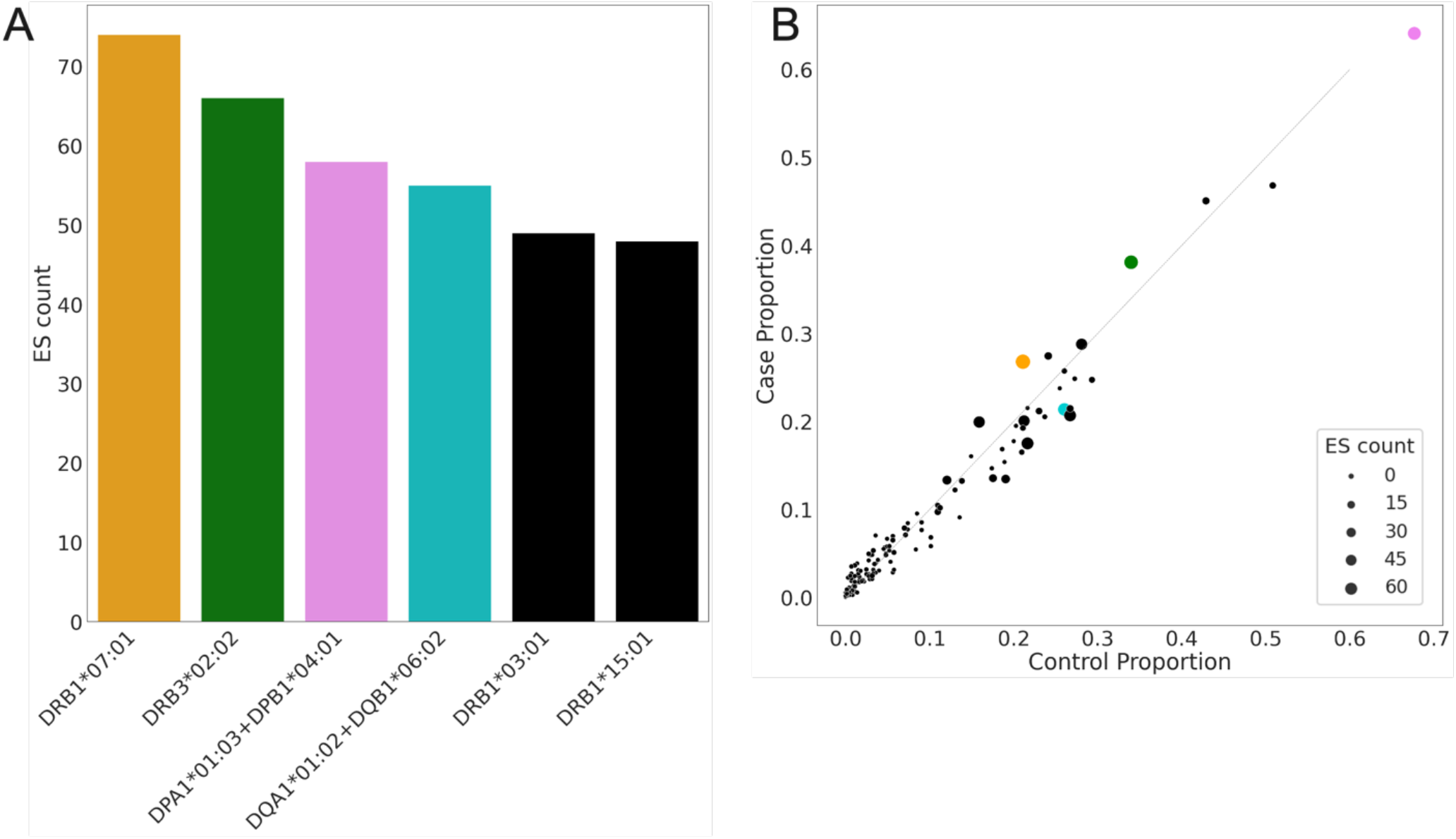
Crohn’s associated ES were statistically associated with presenting HLA heterodimers. A) Counts of ES that could be significantly (p < 0.0001, Fisher’s exact test) associated with an HLA heterodimer. Top 6 alleles are shown. B) Scatterplot of proportion of training controls and cases imputed to have that HLA. Point size is proportional to ES count from (A), colors are consistent with (A). Grey, dotted line indicates equivalent representation of the HLA in cases and controls

### CD ES Breadth is associated with HLA interaction effects

We observed that there were strong associations between disease-specific ES and the complex HLA effects reported. We observed DRB1*07:01 homozygous Crohn’s patients had higher DRB1*07:01 Crohn’s ES breadth than DRB1*07:01 heterozygous patients (Figure 6A). Interestingly, we observed the opposite trend with the DQ6.2/DRB1*15:01 ES – where individual’s heterozygous for those loci had higher Crohn’s ES breadth than individuals homozygous at those loci (Figure 6B). For the subset of DQ6.2/DRB1*15:01 ES, we observed that DQ6.2+/DR15-patients had significantly lower clonal breadth than DQ6.2+/DR15+ and DQ6.2-/DR15+ patients. We also observed that amongst all DQ7.5 heterozygous samples, the subset DQ7.5/DQ4.4 and DQ7.5/DQ2.2 had significantly higher Crohn’s ES clonal breadth (Figure 6C). These results demonstrate how disease specific TCR response is modulated by zygosity and interaction effects at the HLA loci.

**Figure 6:**
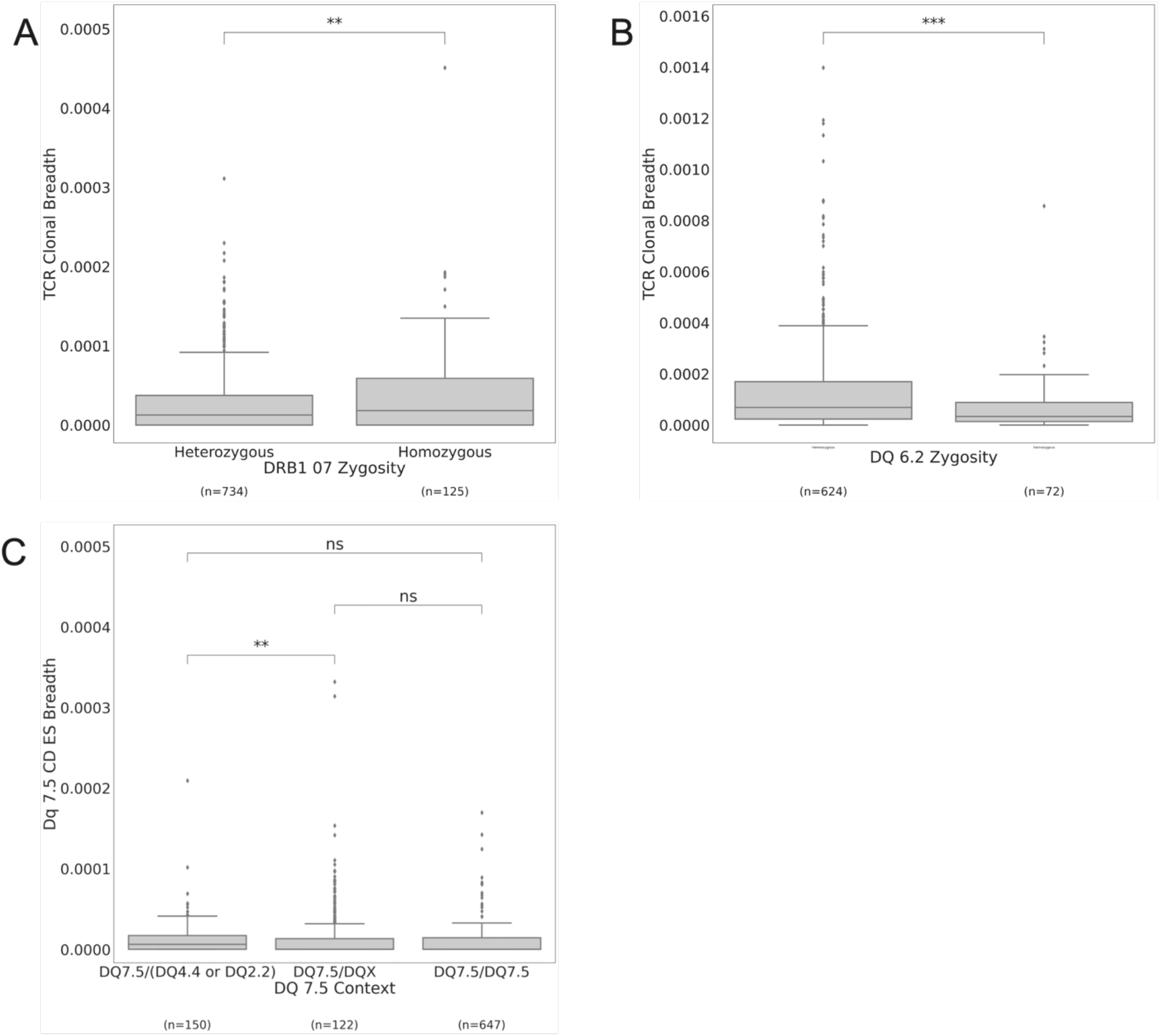
TCR Clonal breadth associations with HLA effects. A) DRB1*07 Homozygous cases had higher disease associated TCR Clonal Breadth in CD (* = p-value <0.05). B) DQ 6.2 Heterozygous cases had higher disease associated TCR Clonal Breadth than DQ6.2 Homozygous cases(*** = p-value <0.0001). C) DQ7.5/DQ2.2 cases had higher disease associated TCR clonal breadth than other DQ7.5 Heterozygous cases in CD (** = p-value <0.001).

### High confidence ES with similar CDR3 sequences associate to the same HLAs

We next investigated if there were subsets of high confidence ES with similar CDR3 amino acid sequences. High confidence ES can be formed into clusters of 1-Hamming distance connected components (Figure 7). Within clusters, ES tend to have the same HLA associations, with two notable exceptions. The cluster with the most included ES (30) and another large cluster of ES with the same TCRB V gene and CDR3 length but different J gene share a strong sequence motif but do not have confident HLA associations for any of their constituent ES (no-HLA, grey cluster, Figure 7). The five clusters with the most ES (>10 each, totaling 92 ES, Figure S10C) are each represented in more than 10% of the CD cases and less than 5% of the healthy controls (Figure S10D). With one exception, no UC ES are within 1-Hamming distance of CD ES (Figure S11). These results imply that the selection of CD ES is antigen-driven, with similar CDR3 motifs associated with a consistent HLA context.

**Figure 7:**
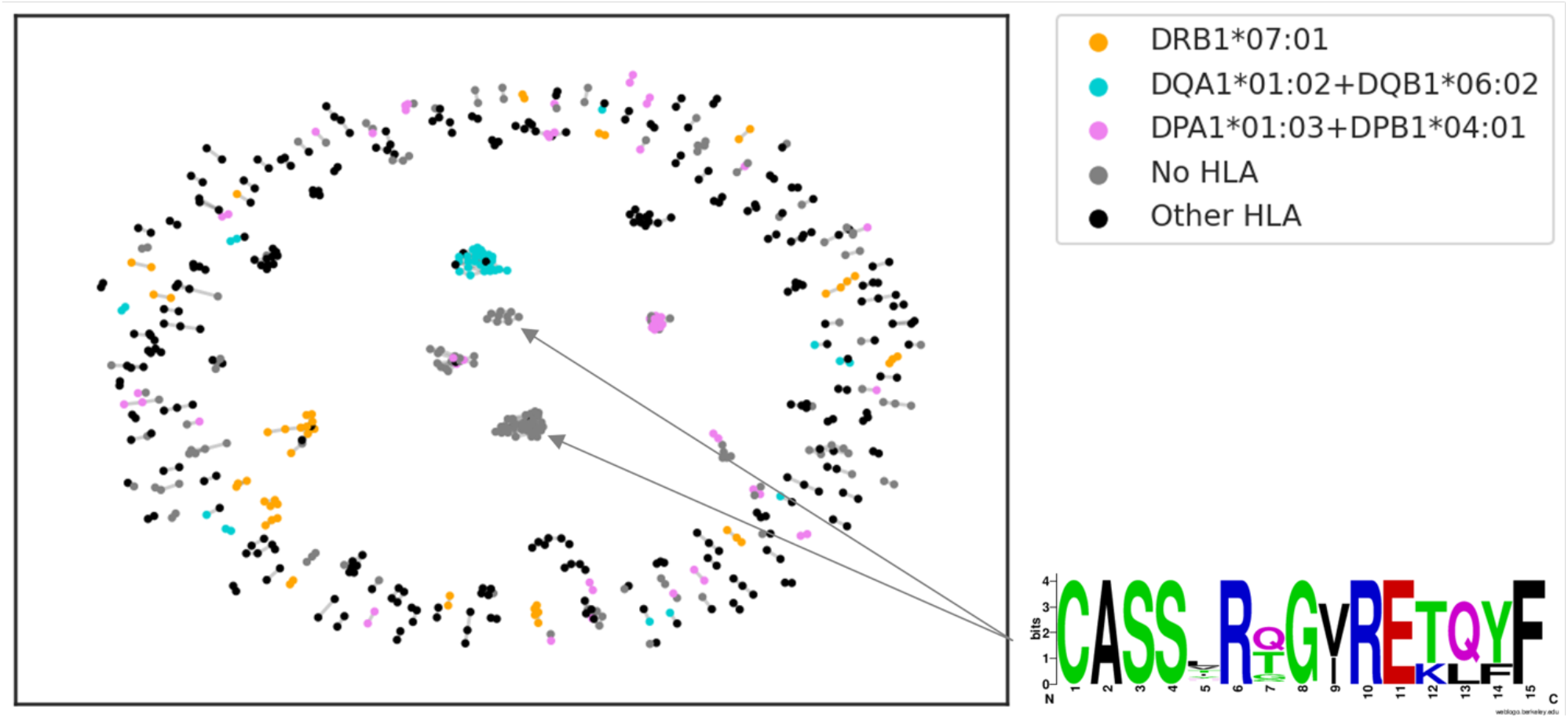
Network graph of high-confidence CD ES, where each edge represents 1-Hamming distance between ES nodes (edge length = similarity metric between ES). Nodes are colored by HLAs that could be significantly associated. Arrows and sequence logo note two clusters that share a CDR3 sequence motif (clusters separated by the J gene portion of the CDR3), where only one ES between the clusters is significantly associated with an HLA.

## Discussion

Previous characterizations of the TCR repertoire in IBD failed to find TCRB sequences specific to IBD. Our success is due, in part, to the size of our case-control study, both in clonotypes per repertoire and in number of repertoires sampled. Our study is the most highly powered study to date to detect more rare clonotypes contributing to IBD in an unbiased manner. An alternative route would be to enrich for IBD-specific T cells using specific antigens; however, to this point it has not been clear that antigens exist which elicit specific and public immune reactions. Although this work does not identify those antigens, it reveals their HLA context and cognate T-cell populations. The TCRs from those T cells are a tractable, blood-based biomarker of IBD, and further work could reveal their targets and role in disease pathology.

While we were able to find specific public TCRs enriched in IBD cases in peripheral blood, the gut is at the center of excessive immune activation in IBD. Indeed, we found the breadth of CD and UC ES were an order of magnitude higher in mucosa samples than in blood, even though the ES were identified in blood originally. Through this lens, the relatively low breadth of UC ES in blood contrasts with the much higher breadth of CD ES in blood, particularly from ileal CD cases. This distinction could be a result of the influence of CCR9 on T-cell localization to the ileum, versus general gut localization through α4β7. The additive effect of fistulas or strictures to ES breadth in blood raises the alternative possibility that exchange with blood is increased in ileal CD due to increased contact with the mesentery. Beyond increased localization and exchange with blood, T-cell populations could be activated outside of gut tissue, as demonstrated in a recent work of showing microbial translocation to the mesentery in ileal CD^23^, which could lead to activation of T-cell populations outside of gut tissues.

Having validated the existence of CD and UC ES across cohorts from different locations and in mucosal samples, our next goal was to characterize them. Working from the paradigm of microbial antigens driving T-cell expansion, we expected high overlap of ES in CD and UC. Instead, we found them to be nearly distinct sets. This distinction may be a reason why a prior effort to identify clonotypes specific to IBD (considering all types together) did not find any significantly enriched sequences^9^. We find that even when limiting to colonic presentations of IBD, where there are shared microbial communities^24^, CD ES do not overlap with UC ES – suggesting that the excessive T-cell response in colonic CD and UC may be driven by distinct antigens.

This divergence in public TCRs between CD and UC is consistent with observations in CD- and UC- specific antibody classes, (Anti-*Saccharomyces cerevisiae* antibodies (ASCA) and anti-neutrophil cytoplasmic antibodies (ANCA), respectively), which are employed in developing algorithms for serologic diagnostics to distinguish these diseases^25^. The apparent T- and B-cell repertoire distinction between CD and UC contrasts with the risk spectrum of IBD described in a highly powered genetic association study^26^ but mostly agrees with the risk conferred by individual HLA alleles^27^.

It is increasingly clear that despite overlapping genetic risk, CD and UC involve distinct patterns in the adaptive immune repertoire. By comparing the CD and UC ES against public data, we determined that some CD ES are likely activated by yeast antigens. Further work is required to determine the proportion of the CD-specific T cell response attributable to yeast, and to identify remaining activators in CD and UC.

The higher numbers of CD samples and CD ES, compared to those of UC, gave us increased power to determine additional characteristics of CD ES. The first of these was the HLA each CD ES is most likely to bind. Although our HLA models only cover 145 HLA alleles, they are the 145 HLA alleles that are most common in the populations from the regions included in this study (Europe and the United States). Because TCRs binding rarer HLAs are unlikely to be public in our sequence discovery set, they are not likely to be identified as ES. The fraction of CD ES associated to each HLA then reflects both the HLA’s involvement in disease and its prevalence in the population.

The CD ES are primarily associated to class II HLAs, which aligns with findings from HLA-focused genetic studies in IBD^4,5^. Yet, regarding individual HLA alleles, the agreement is variable. Although DRB1*01:03 has been strongly associated to colonic CD and UC^5,26^, we found only a single CD ES associated to it. This allele was present in fewer than 5% of our IBD samples, which gave us low statistical power to identify TCRs binding it as ES. On the other hand, DRB1*07:01 has been associated with increased risk for ileal CD^5,26^ and is also the HLA with the most ES associated in our study. The DR15 haplotype (including DRB1*15:01 (DR15), DQA1*01:02+DQB1*06:02 (DQ6.2), and DRB5*01:01) has been associated to increased UC risk, but not increased CD risk^4,5^, which is also observed in our study. In fact, DR15 and DQ6.2 both show a protective effect in CD. Comparably, DRB3:02:02 and DPA1*01:03+DPB1*04:01 (DP401.1) each had >50 ES associated to them but have not been linked to increased risk for any IBD subgroup. Previous studies^5^ that used sequencing or probe-based approach to identify interaction effects may have failed to resolve complex interactions between the DQ and DP heterodimers. These results demonstrate the feasibility and advantages of using TCRβ repertoires to identify complex HLA interactions in the HLA loci contributing to disease risk. Taken together, these associations suggest that class II HLAs drive a specific T-cell response by presenting similar peptides in subgroup of patients with the same haplotypes. The inflammatory or protective functions of the T cells encoding ES remains to be elucidated. We posit that the methods developed and applied here can be used in other disease areas to understand how complex HLA effects shape disease associated TCR response.

We found that, although sequence similarity was not used to associate ES to HLAs, groups of CD ES connected by 1-Hamming distance connections (in the CDR3 region) generally had a single HLA association. This conservation implies that the TCRB CDR3 region is either mediating the HLA interaction, or interaction with a restricted set of peptides presented by a single HLA. One subpopulation of CD ES does not associate to any of our imputed HLAs, despite its presence in nearly 20% of CD cases. This population is similar to the recently described CAIT cells in that they have a strong CDR3 motif, are specific to CD, and do not bind conventional HLAs^10^. We cannot rule out that these TCRB sequences are paired with CAIT TRA sequences, but they do not match any of the published CAIT TCRB sequences. A recently developed tool for identifying MAIT cells from TCRB sequence data^28^ predicted these were also unlikely to be MAIT cells, though we cannot rule out the possibility. Even if they are not CAIT, they could still be part of the same type II NKT population, potentially reinforcing the importance of unconventional T cells in CD.

This study has revealed common T-cell receptor beta sequences specific to CD and UC, but further work will be required to determine their role in disease. It is not immediately apparent what set of antigens could be provoking this specific yet widespread response, since there are not particular pathogens or autoantigens known to be unique to either CD or UC. ASCA and ANCA suggest potential search spaces for antigens,. Separate from their targets, it is not clear if ES participate to the pathophysiology of IBD or appear as a consequence of the general increased immune response in the intestinal mucosa. While these two processes are not mutually exclusive, the persistence of specific clones over time associated with disease activity advocate for the former. At the very least, ES are a potential new disease marker in IBD but further characterization, especially their patterns throughout disease courses, could also lead to new insights into their involvement in disease mechanisms.

## Acknowledgements

The results published here are in part based on data obtained from the IBD Plexus program of the Crohn’s & Colitis Foundation.

## References

1. Lewis, J. D. et al. Incidence, Prevalence, and Racial and Ethnic Distribution of Inflammatory Bowel Disease in the United States. Gastroenterology 165, 1197-1205.e2 (2023).

2. Burisch, J., Jess, T., Martinato, M., Lakatos, P. L. & ECCO -EpiCom. The burden of inflammatory bowel disease in Europe. J Crohns Colitis 7, 322–37 (2013).

3. Guan, Q. A Comprehensive Review and Update on the Pathogenesis of Inflammatory Bowel Disease. J Immunol Res 2019, 7247238 (2019).

4. Degenhardt, F. et al. Transethnic analysis of the human leukocyte antigen region for ulcerative colitis reveals not only shared but also ethnicity-specific disease associations. Hum Mol Genet 30, 356–369 (2021).

5. Goyette, P. et al. High-density mapping of the MHC identifies a shared role for HLA-DRB1*01:03 in inflammatory bowel diseases and heterozygous advantage in ulcerative colitis. Nat Genet 47, 172–9 (2015).

6. Ahmad, T.Marshall, S.-E. & Jewell, D. Genetics of inflammatory bowel disease: the role of the HLA complex. World J Gastroenterol 12, 3628–35 (2006).

7. Gittelman, R. M. et al. Longitudinal analysis of T cell receptor repertoires reveals shared patterns of antigen-specific response to SARS-CoV-2 infection. JCI Insight 7, (2022).

8. Greissl, J. et al. Immunosequencing of the T-Cell Receptor Repertoire Reveals Signatures Specific for Identification and Characterization of Early Lyme Disease. medRxiv 2021.07.30.21261353 (2022) doi:10.1101/2021.07.30.21261353.

9. Rosati, E. et al. Identification of Disease-associated Traits and Clonotypes in the T Cell Receptor Repertoire of Monozygotic Twins Affected by Inflammatory Bowel Diseases. J Crohns Colitis 14, 778–790 (2020).

10. Rosati, E. et al. A novel unconventional T cell population enriched in Crohn’s disease. Gut 71, 2194–2204 (2022).

11. Uchida, A. M. et al. Escherichiacoli-Specific CD4+ T Cells Have Public T-Cell Receptors and Low Interleukin 10 Production in Crohn’s Disease. Cell Mol Gastroenterol Hepatol 10, 507–526 (2020).

12. Li, J. et al. Profiles of Lamina Propria T Helper Cell Subsets Discriminate Between Ulcerative Colitis and Crohn’s Disease. Inflamm Bowel Dis 22, 1779–92 (2016).

13. Bouma, G. & Strober, W. The immunological and genetic basis of inflammatory bowel disease. Nat Rev Immunol 3, 521–33 (2003).

14. Hong, S. N. et al. Reduced diversity of intestinal T-cell receptor repertoire in patients with Crohn’s disease. Front Cell Infect Microbiol 12, 932373 (2022).

15. Allez, M. et al. T cell clonal expansions in ileal Crohn’s disease are associated with smoking behaviour and postoperative recurrence. Gut 68, 1961–1970 (2019).

16. Carlson, C. S. et al. Using synthetic templates to design an unbiased multiplex PCR assay. Nat Commun 4, 2680 (2013).

17. Robins, H. et al. Ultra-sensitive detection of rare T cell clones. J Immunol Methods 375, 14–9 (2012).

18. Pruessmann, W. et al. Molecular analysis of primary melanoma T cells identifies patients at risk for metastatic recurrence. Nat Cancer 1, 197–209 (2020).

19. Zahid, H. J. et al. Large-scale statistical mapping of T-cell receptor β sequences to Human Leukocyte Antigens. bioRxiv 2024.04.01.587617 (2024) doi:10.1101/2024.04.01.587617.

20. Emerson, R. O. et al. Immunosequencing identifies signatures of cytomegalovirus exposure history and HLA-mediated effects on the T cell repertoire. Nat Genet 49, 659–665 (2017).

21. Stokkers, P. C., Reitsma, P. H., Tytgat, G. N. & van Deventer, S. J. HLA-DR and -DQ phenotypes in inflammatory bowel disease: a meta-analysis. Gut 45, 395–401 (1999).

22. Martini, G. R. et al. Selection of cross-reactive T cells by commensal and food-derived yeasts drives cytotoxic TH1 cell responses in Crohn’s disease. Nat Med 29, 2602–2614 (2023).

23. Ha, C. W. Y. et al. Translocation of Viable Gut Microbiota to Mesenteric Adipose Drives Formation of Creeping Fat in Humans. Cell 183, 666-683.e17 (2020).

24. Lloyd-Price, J. et al. Multi-omics of the gut microbial ecosystem in inflammatory bowel diseases. Nature 569, 655–662 (2019).

25. Sokollik, C. et al. Machine Learning in Antibody Diagnostics for Inflammatory Bowel Disease Subtype Classification. Diagnostics (Basel) 13, (2023).

26. Cleynen, I. et al. Inherited determinants of Crohn’s disease and ulcerative colitis phenotypes: a genetic association study. Lancet 387, 156–67 (2016).

27. Ye, B. D. & McGovern, D. P. B. Genetic variation in IBD: progress, clues to pathogenesis and possible clinical utility. Expert Rev Clin Immunol 12, 1091–107 (2016).

28. ElAbd, H. et al. Seq2MAIT: A Novel Deep Learning Framework for Identifying Mucosal Associated Invariant T (MAIT) Cells. bioRxiv 2024.03.12.584395 (2024) doi:10.1101/2024.03.12.584395.

