## Supplementary Material for "Antigen-driven expansion of public clonal T cell populations in inflammatory bowel diseases"

### Supplementary Information

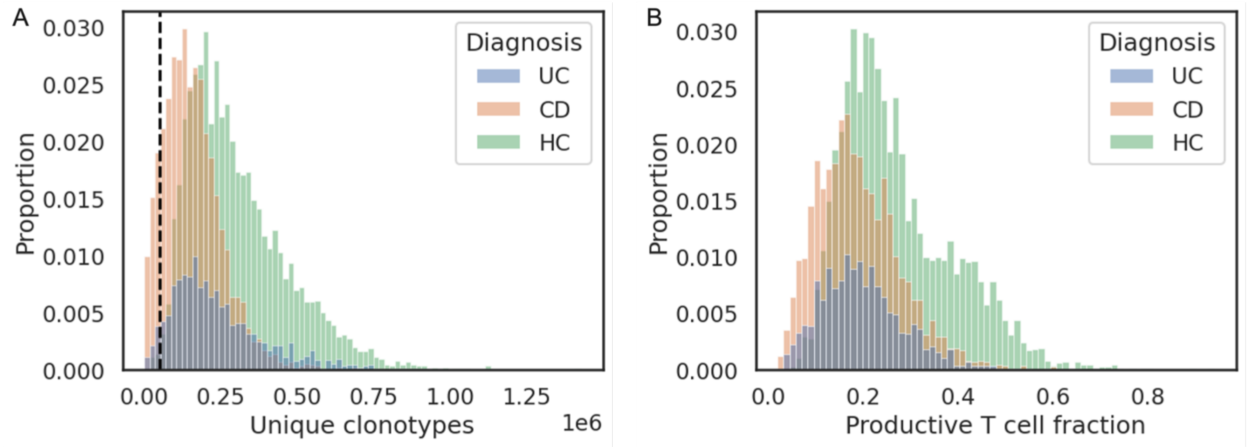


Figure S1: Distributions of unique clonotypes sequenced (A) and productive T-cell fraction (B) for CD, UC, and healthy controls. Black dashed line in panel (A) undicates lower QC threshold for analyses.

| | | | | Singleton Breadth | | | | | | | Simpson Clonality | | | | | | | Clonotype Count | | | | | | |
|---|---|---|---|---|---|---|---|---|---|---|---|---|---|---|---|---|---|---|---|---|---|---|---|---|
| Dx | CMV | age | count | mean | std | min | 25% | 50% | 75% | max | mean | std | min | 25% | 50% | 75% | max | mean | std | min | 25% | 50% | 75% | max |
| CD | Neg | 20-39 | 1157 | 0.92 | 0.03 | 0.76 | 0.91 | 0.93 | 0.95 | 0.99 | 0.02 | 0.02 | 0.00 | 0.01 | 0.01 | 0.02 | 0.33 | 40628 | 4944 | 18717 | 38101 | 41512 | 44292 | 48803 |
| | | 40-59 | 637 | 0.91 | 0.04 | 0.75 | 0.89 | 0.92 | 0.94 | 0.97 | 0.03 | 0.03 | 0.01 | 0.01 | 0.02 | 0.03 | 0.35 | 38798 | 5420 | 15898 | 35973 | 39658 | 42866 | 47727 |
| | | 60-79 | 172 | 0.89 | 0.04 | 0.73 | 0.87 | 0.90 | 0.92 | 0.97 | 0.03 | 0.04 | 0.01 | 0.01 | 0.02 | 0.04 | 0.35 | 36807 | 5965 | 10180 | 33470 | 37391 | 41348 | 46822 |
| | Pos | 20-39 | 535 | 0.92 | 0.03 | 0.80 | 0.90 | 0.93 | 0.94 | 0.98 | 0.06 | 0.05 | 0.01 | 0.03 | 0.04 | 0.07 | 0.42 | 34244 | 7162 | 7384 | 30282 | 35535 | 39447 | 47806 |
| | | 40-59 | 397 | 0.91 | 0.04 | 0.79 | 0.88 | 0.91 | 0.93 | 0.98 | 0.07 | 0.05 | 0.01 | 0.03 | 0.05 | 0.08 | 0.46 | 31887 | 7458 | 4176 | 27172 | 33308 | 37218 | 46349 |
| | | 60-79 | 158 | 0.90 | 0.03 | 0.80 | 0.88 | 0.90 | 0.92 | 0.97 | 0.09 | 0.07 | 0.01 | 0.04 | 0.08 | 0.12 | 0.47 | 28664 | 8056 | 9230 | 22650 | 29094 | 34932 | 43710 |
| Control | Neg | 20-39 | 621 | 0.94 | 0.02 | 0.83 | 0.93 | 0.94 | 0.96 | 0.99 | 0.02 | 0.01 | 0.00 | 0.01 | 0.01 | 0.02 | 0.20 | 42551 | 3481 | 23290 | 40621 | 43009 | 45104 | 48577 |
| | | 40-59 | 430 | 0.92 | 0.03 | 0.81 | 0.91 | 0.93 | 0.95 | 0.98 | 0.02 | 0.02 | 0.01 | 0.01 | 0.02 | 0.03 | 0.16 | 39667 | 4514 | 20437 | 37337 | 40414 | 42806 | 47966 |
| | | 60-79 | 137 | 0.91 | 0.03 | 0.80 | 0.89 | 0.91 | 0.94 | 0.97 | 0.03 | 0.04 | 0.00 | 0.01 | 0.02 | 0.04 | 0.24 | 38265 | 5228 | 22642 | 35048 | 38744 | 42429 | 47891 |
| | Pos | 20-39 | 650 | 0.93 | 0.02 | 0.85 | 0.92 | 0.93 | 0.95 | 0.98 | 0.04 | 0.03 | 0.00 | 0.02 | 0.03 | 0.05 | 0.39 | 38230 | 5036 | 16995 | 35140 | 38971 | 41873 | 48032 |
| | | 40-59 | 518 | 0.92 | 0.03 | 0.75 | 0.90 | 0.92 | 0.94 | 0.98 | 0.05 | 0.04 | 0.01 | 0.02 | 0.04 | 0.07 | 0.22 | 35279 | 5253 | 17918 | 31873 | 35526 | 39264 | 46293 |
| | | 60-79 | 195 | 0.91 | 0.03 | 0.80 | 0.90 | 0.91 | 0.93 | 0.98 | 0.08 | 0.06 | 0.01 | 0.04 | 0.06 | 0.10 | 0.35 | 32058 | 6807 | 7976 | 27767 | 32240 | 37056 | 46237 |
| UC | Neg | 20-39 | 429 | 0.93 | 0.03 | 0.79 | 0.92 | 0.94 | 0.95 | 0.99 | 0.02 | 0.02 | 0.00 | 0.01 | 0.01 | 0.02 | 0.24 | 41801 | 3932 | 25082 | 39774 | 42506 | 44618 | 48882 |
| | | 40-59 | 317 | 0.92 | 0.03 | 0.82 | 0.90 | 0.92 | 0.94 | 0.98 | 0.02 | 0.03 | 0.00 | 0.01 | 0.02 | 0.03 | 0.35 | 40107 | 4648 | 19100 | 37550 | 40488 | 43621 | 48308 |
| | | 60-79 | 92 | 0.91 | 0.03 | 0.83 | 0.89 | 0.91 | 0.93 | 0.97 | 0.02 | 0.02 | 0.01 | 0.01 | 0.02 | 0.03 | 0.11 | 38879 | 4466 | 21671 | 36297 | 39107 | 42280 | 46859 |
| | Pos | 20-39 | 207 | 0.93 | 0.03 | 0.84 | 0.92 | 0.93 | 0.95 | 0.98 | 0.06 | 0.04 | 0.01 | 0.03 | 0.05 | 0.08 | 0.33 | 34123 | 6886 | 13389 | 29837 | 35293 | 39089 | 45826 |
| | | 40-59 | 190 | 0.92 | 0.04 | 0.62 | 0.90 | 0.92 | 0.94 | 0.98 | 0.07 | 0.06 | 0.01 | 0.04 | 0.06 | 0.10 | 0.38 | 31279 | 7631 | 3720 | 25471 | 31974 | 36940 | 45133 |
| | | 60-79 | 109 | 0.90 | 0.04 | 0.75 | 0.89 | 0.91 | 0.93 | 0.96 | 0.09 | 0.06 | 0.01 | 0.05 | 0.08 | 0.13 | 0.28 | 28574 | 8707 | 6021 | 22316 | 29787 | 34862 | 47303 |

Table S1: Distribution metrics of samples represented by each boxplot in Figure 1A-1C based on diagnosis, age bin, and CMV status.

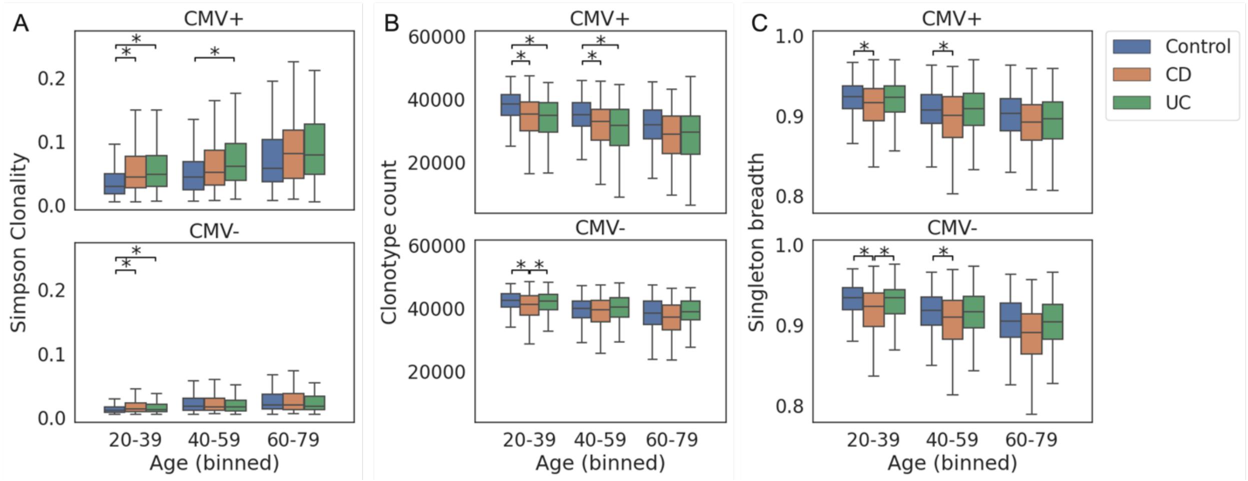


Supplementary Figure S2: After downsampling to 50,000 total read templates and applying an empiric downsampling correction, in all samples from individuals with attached age data, we assessed A) Simpson clonality, B) clonotype count, and C) singleton breadth for CD and UC cases and healthy controls, across age bins and CMV status. All pairwise comparisons within an age and CMV bin were assessed with a Kolmogorov–Smirnov test for equivalence of the distributions. All comparisons with $p < 0.05$ are indicated with a '*', though in this case all marked comparisons have $p < 2x10^{-4}$. Each boxplot represents the distribution of normalized clonotype counts in that bin across samples. Boxes cover the first and third quartile of the distribution (with the median indicated within the box), while whiskers show the remainder of the distribution up to the interquartile range. Outliers were excluded to better visualize the central quantiles of the distribution. Sample counts are the same as in Figure 1D.

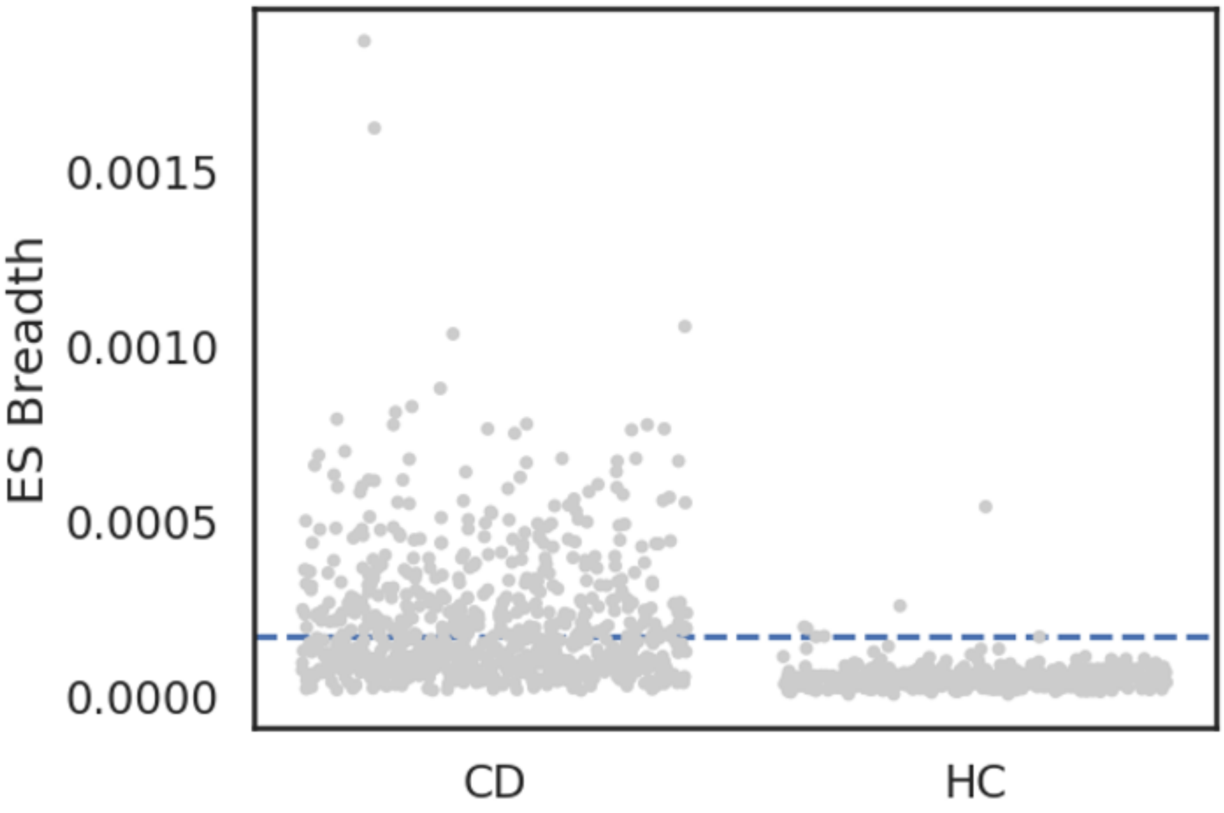


Figure S3: Strip plot of CD ES breadth for each CD (N=758) and HC (N=452) sample in the validation set. CD ES breadth is significantly higher ($p < 1x10^{-100}$).

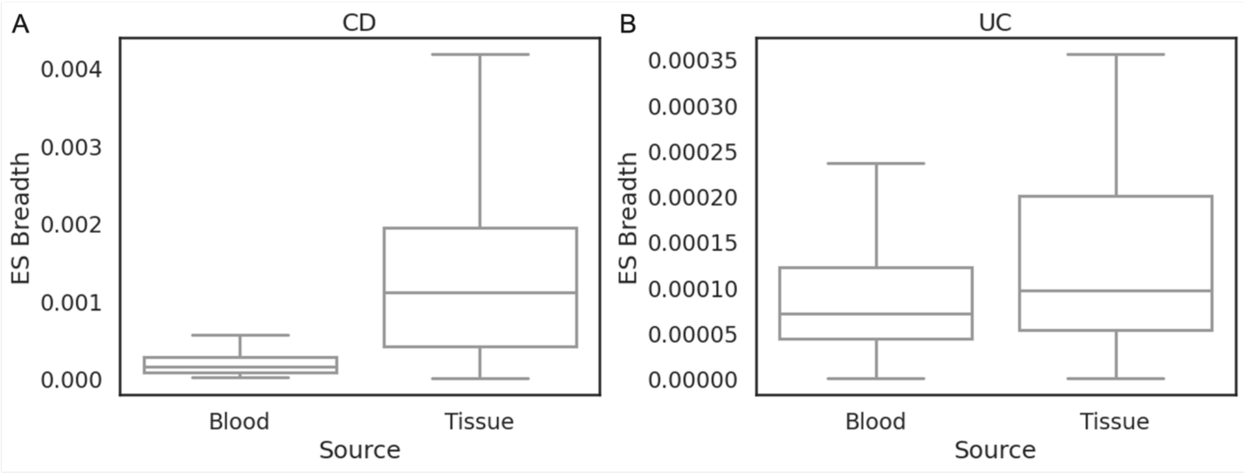


Figure S4: ES breadth differences between blood and tissue for (A) CD ES in CD cases and (B) UC ES in UC cases.

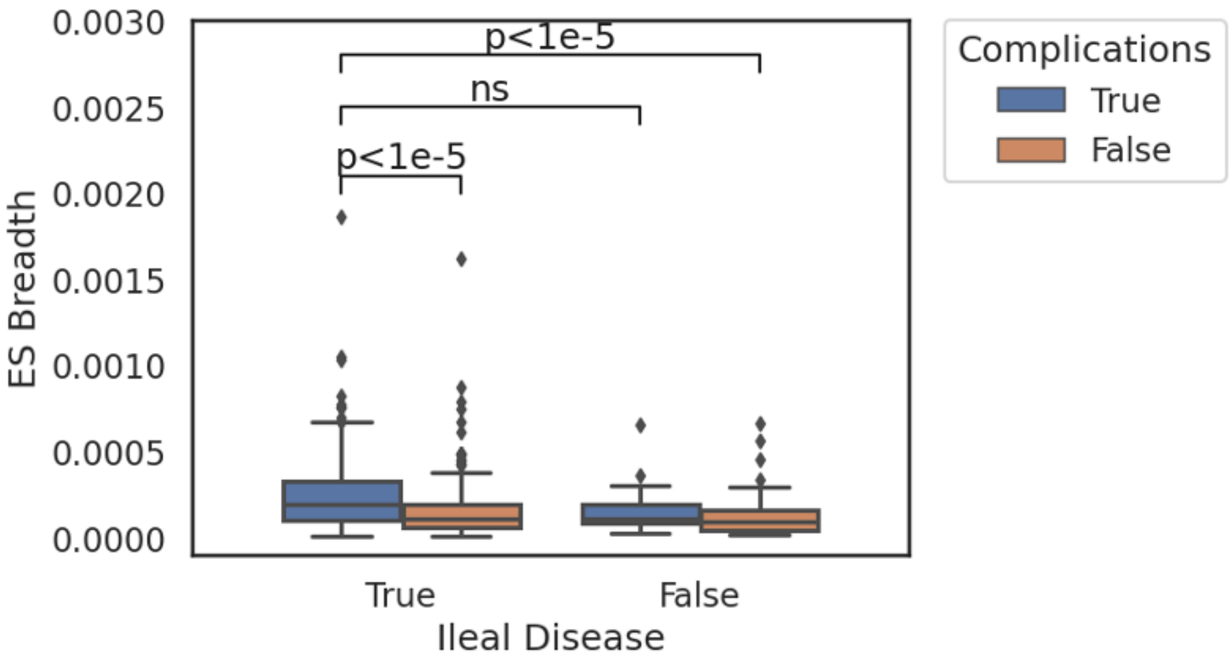


Figure S5: CD ES breadth is significantly higher in cases with ileal involvement and stricturing and/or fistulizing behavior (complications, N=163) than in cases with only colonic involvement without complications (N=45) or cases with ileal involvement without complications (N=149). Cases with complications and only colonic involvement (N=16) did not have significantly different breadth from other categories, potentially due to low sample numbers.

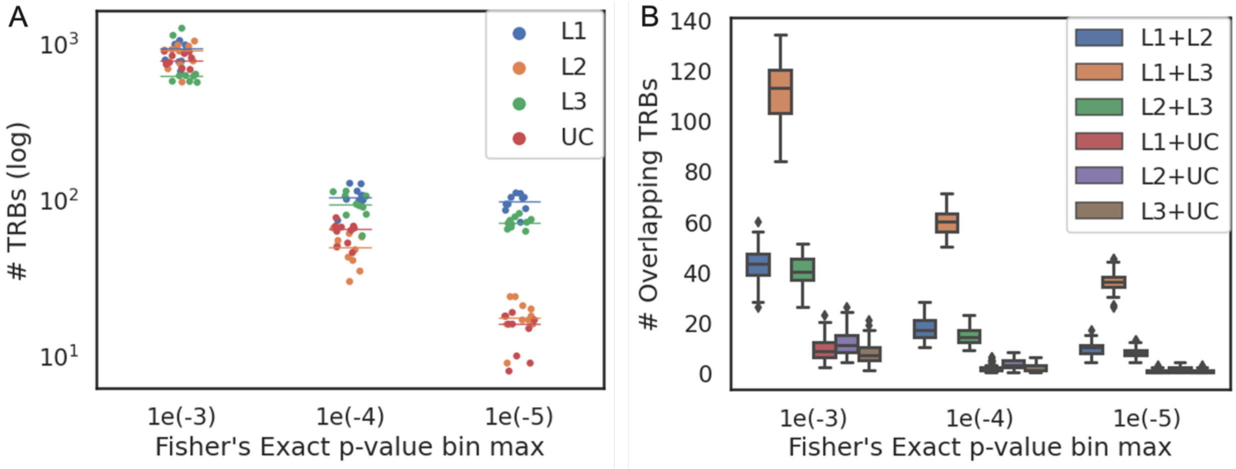


Figure S6: Identification of specific TRB sequences enriched in CD and UC. A) Count of TRB sequences in repeat experiments. Within each p-value threshold, each dot represents an experiment comparing 500 randomly selected disease repertoires to 2,607 control repertoires. 10 experiments were performed for each disease. Median values shown as a horizontal line. B) Box and whisker plot of specific TRB sequences that were significant in multiple IBD subtypes. All replicates for each IBD subtype were compared pairwise, resulting in 100 comparisons for each box. L1=ileal CD, L2=colonic CD, L3=ileocolonic CD.

| | L1+L2 | L1+L3 | L2+L3 | L1+UC | L2+UC |
|---|---|---|---|---|---|
| **L1+L3** | $3.1x10^{-33}$ | | | | |
| **L2+L3** | 0.021 | $2.7x10^{-33}$ | | | |
| **L1+UC** | $4.8x10^{-34}$ | $5.1x10^{-34}$ | $4.1x10^{-34}$ | | |
| **L2+UC** | $2.1x10^{-33}$ | $1.5x10^{-33}$ | $1.5x10^{-33}$ | 0.025 | |
| **L3+UC** | $7.8x10^{-35}$ | $8.3x10^{-35}$ | $6.7x10^{-35}$ | 0.28 | $1.4x10^{-5}$ |

Table S2: Table of p-values from the Mann Whitney U test for pairwise comparison total overlapping TCR sequences, after Bonferroni correction for testing 15 hypotheses. L1=ileal CD, L2=colonic CD L3=ileocolonic CD.

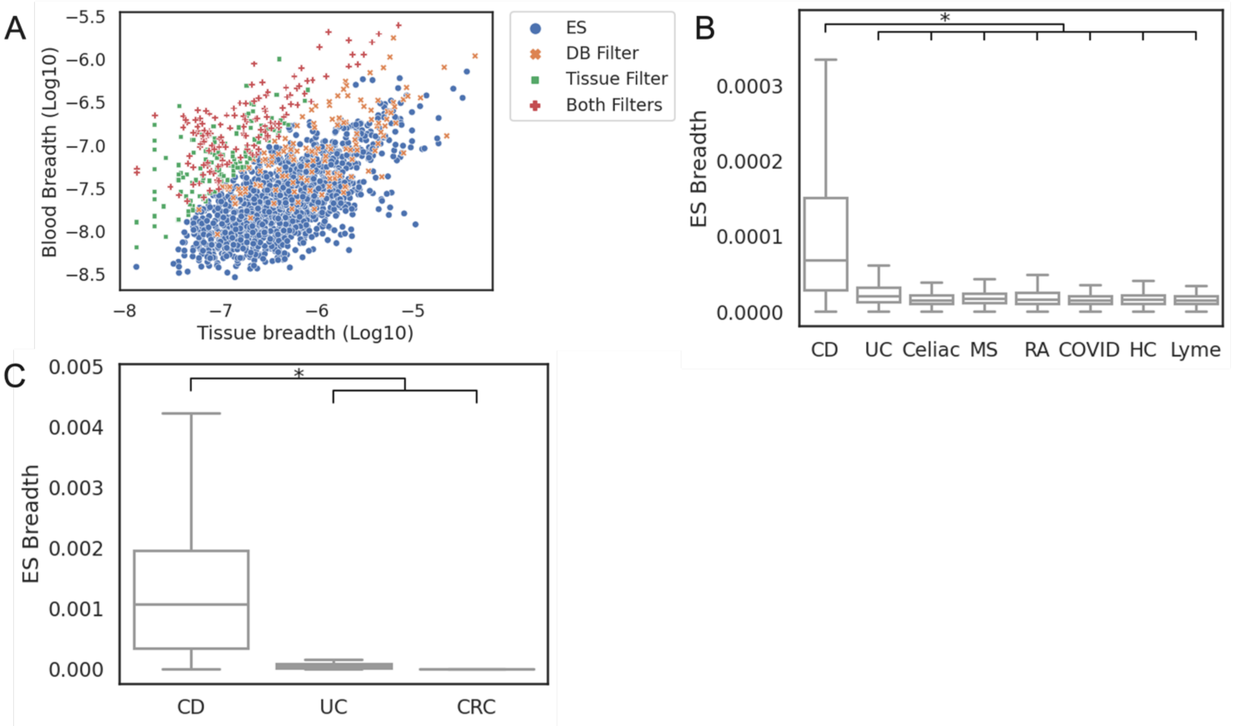


Figure S7: Distribution of sequence breadth of Crohn's associated ES in training set blood samples and matched tissue samples (A), breadth of filtered ES in holdout blood samples (B), and breadth of filtered ES in holdout tissue samples (C). Boxplots show first and third quartile values of the breadth range, with the median marked within the box. Whiskers extend to furthest point within 1.5 times the interquartile range. Points outside of the whiskers were excluded for visualization purposes. CD = Crohn's disease, UC = ulcerative colitis, HC = healthy control, MS = multiple sclerosis, RA = rheumatoid arthritis, CRC = colorectal cancer.

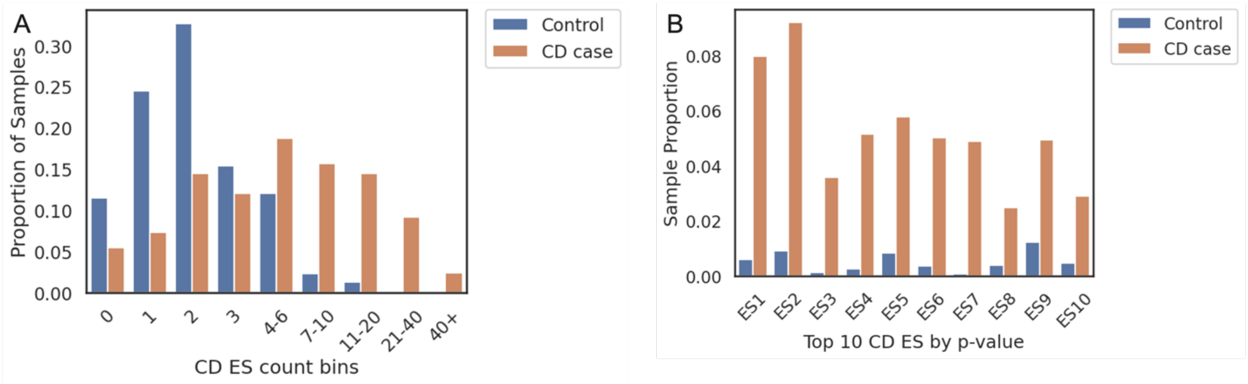


Figure S8: A) Distribution of CD ES per sample for healthy controls and CD cases. Y-axis is proportion of samples from either group who have the ES in the bin on the x-axis. B) Proportion of healthy control or CD case individuals who have the 10 ES most specific for CD by FET p-value.

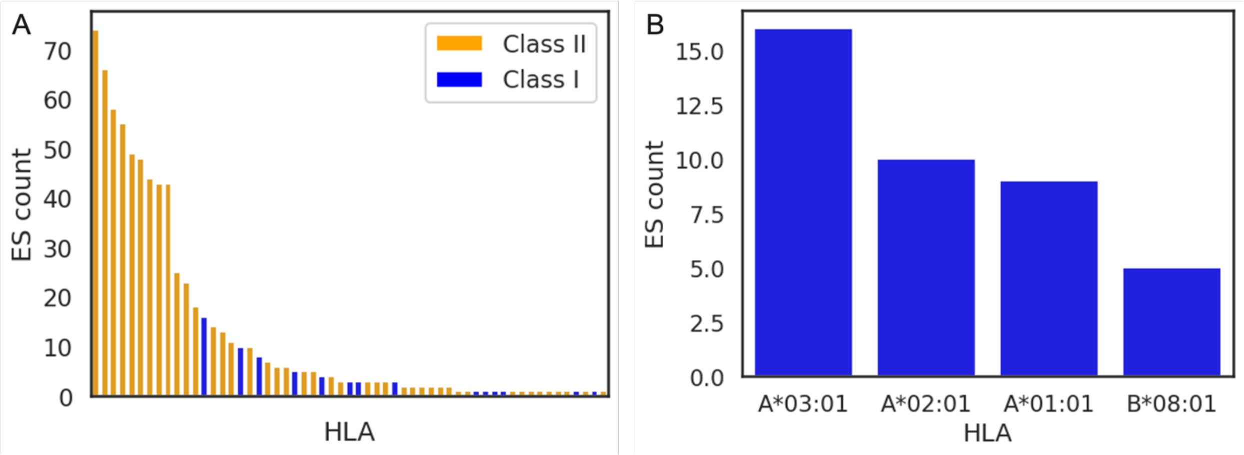


Figure S9: A) Distribution of associated ES count by HLA for all repertoire-imputed HLAs with at least one associated ES. B) Distribution of associated ES count by HLA for the 5 class I HLAs with the most associated ES.

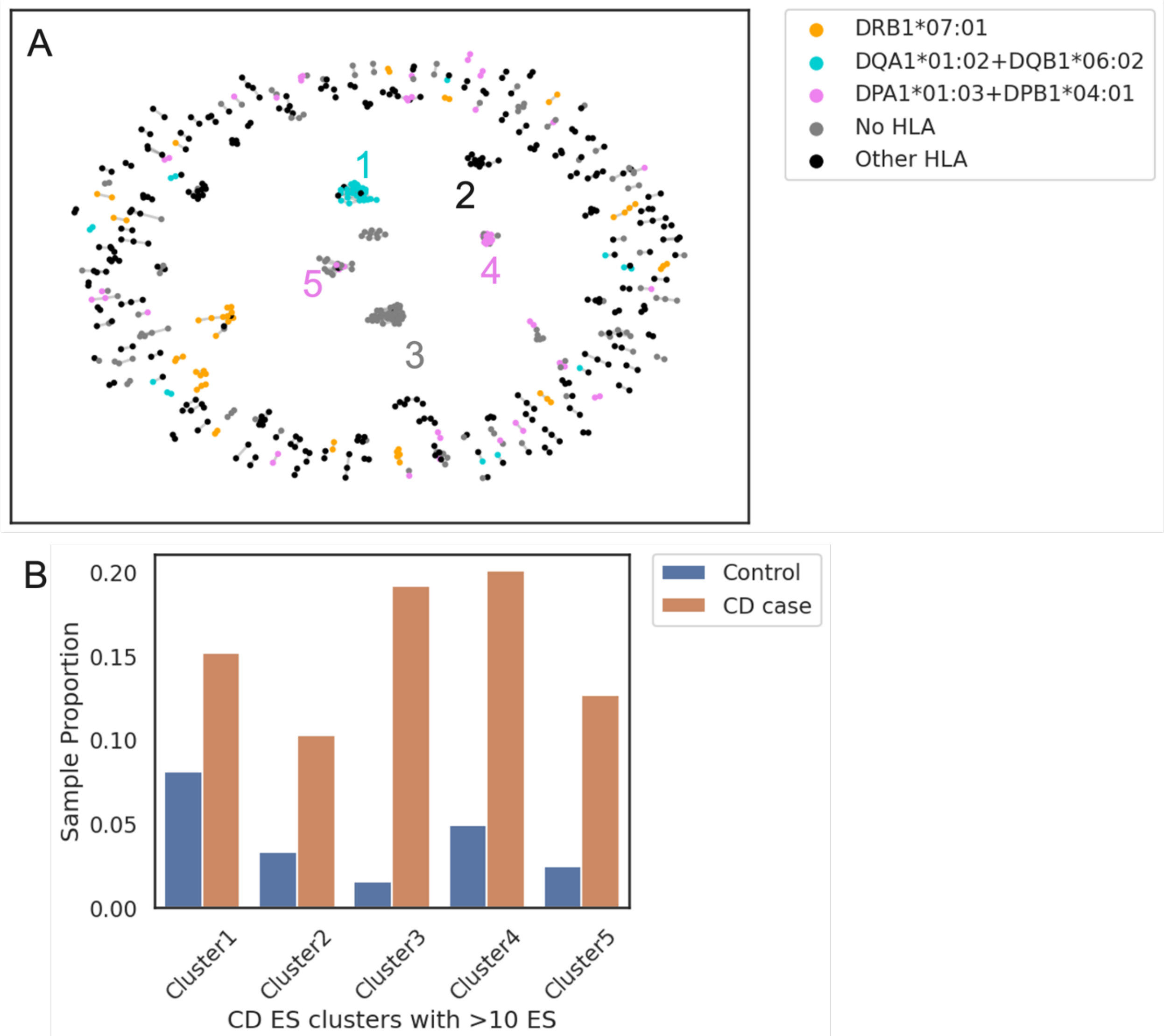


Figure S10: A) Cluster diagram as in Figure 7 with the five largest clusters labeled. B) Proportion of healthy control or CD case individuals who have at least one ES from the five clusters labeled in part A.

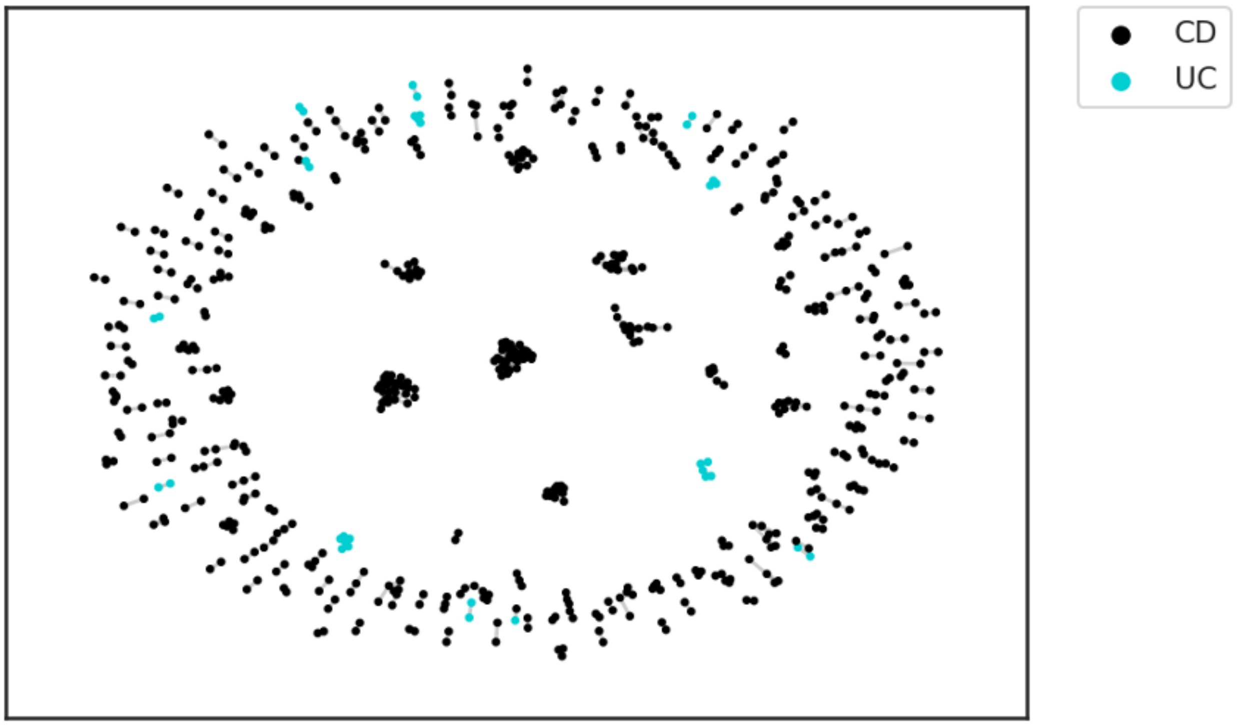


Figure S11: 1-Hamming distance cluster diagram of CD and UC ES together. Clustering and visualization as in Figure 7.
